# Quantitative Modeling of Insulin Signaling Reveals Mechanisms of Insulin Resistance in Obese Mice

**DOI:** 10.1101/2025.10.15.682504

**Authors:** So Morishita, Shinsuke Uda, Hiroyuki Kubota

## Abstract

Insulin resistance is a deficiency in insulin-mediated regulation of glucose metabolism and is considered a cause of multiple diseases such as diabetes and hypertension. The insulin signaling pathway’s contribution to insulin resistance remains incompletely understood. Insulin receptor (IR) mutations can cause insulin resistance; however, reduced IR expression in obesity may still permit sufficient signaling due to the characteristics of IR, known as the spare receptor hypothesis. Selective insulin resistance, in which glucose metabolism is impaired while other pathways remain intact, further complicates insulin resistance. To elucidate these mechanisms, we developed mathematical models of the insulin signaling pathway in mice with diet-induced obesity and found that changes in protein expression levels can largely account for signal attenuation. Insulin resistance was characterized using two indices: sensitivity shift and responsiveness shift. We found a trade-off between the two, whereby loss of responsiveness from upstream decreases was compensated for by decreases in downstream sensitivity. The excess of receptors conferred robustness to receptor decreases via the signaling pathway. Incoherent-feedforward-loop regulation maintained some signaling despite attenuation, potentially explaining selective insulin resistance. These findings provide deeper insights into insulin resistance and quantitative mechanisms underlying signaling pathway regulation.

## 1 Introduction

Insulin resistance refers to impaired glucose metabolism mediated by insulin and is considered to cause various diseases, including diabetes and cardiovascular dysfunction[1,2]. Previous studies have reported mutations and abnormalities in molecules of the insulin signaling pathway, including the insulin receptor (IR), which are associated with insulin resistance[3,4]. Additionally, obesity is a major cause of insulin resistance and has been investigated by numerous studies[5]. Decreased levels of expression of IR and IRS2 in these studies further support this notion[6–11]. On the other hand, a study using adipocytes demonstrated that a full biological response can still be elicited even with only 2.4% of the IR present[12]. Furthermore, Akt phosphorylation, as well as glucose and insulin tolerance, is normal in mice with heterozygous IR deficiency[13]. These findings suggest that a decrease in the IR alone may not be the primary cause of insulin resistance. The phenomenon, in which a similar response is observed despite a decrease in the IR, is explained by the “spare receptor hypothesis”[14]. This is a characteristic common to many receptors, not just the IR, in which the total number of receptors considerably exceeds the number of ligand-binding receptors. The significance of the spare receptors is that they confer robustness against receptor loss due to disease and ensure linearity between the receptor and ligands[14,15]. In other words, even a small quantity of IRs can elicit a sufficient insulin response since only a small number are needed to bind to insulin to achieve a sufficient response compared to the total number of IRs present. While this hypothesis is intuitively comprehensible, its underlying mechanisms remain poorly understood. Additionally, one of the unresolved issues in insulin resistance is the clarification of selective insulin resistance, which is the phenomenon in which insulin-mediated responses, such as glucose metabolism, are impaired, while reactions such as lipid synthesis and protein synthesis remain intact[16,17]. Such selective impairment has also been observed in the insulin signaling pathway molecules, suggesting that some pathways are affected while others are not[2,16]. However, the mechanism of selective specificity remains unclear. Thus, there are still many unresolved issues regarding the significance and mechanisms of the insulin signaling pathway in insulin resistance.

Obesity is a major cause of insulin resistance. Diet-induced obese mice are considered a model that reflects human obesity[18]. In this study, we focused on changes in the dynamics of the insulin signaling pathways associated with insulin resistance. We used mice with high-fat diet (HFD)-induced obesity and developed a mathematical model to understand the mechanisms by which insulin resistance develops. By comparing mathematical models developed under pre-obesity induction (CD5w), post-obesity induction (HFD14w), or non-obese (CD14w) conditions, we demonstrated that changes in protein expression alone can account for alterations in information transmission in the insulin signaling pathway. Analysis using our mathematical models revealed that the molecules examined in this study can be classified into two groups based on their network structure: (1) feedforward (FF) regulation, which includes IR, Akt, GSK3β, FoxO1, and ACLY; and (2) incoherent-feedforward-loop (IFFL) regulation, which includes S6K and 4E-BP. The characteristics of FF were then examined using a toy model and applied to the insulin signaling pathway model we developed. To understand insulin resistance, two indices have been proposed: a sensitivity shift, in which the dose–response curve shifts to the right; and a responsiveness shift, in which the maximum response is reduced[2,19,20]. We applied these two indices to investigate the insulin signaling pathway in the context of insulin resistance. In our analysis, the sensitivity and responsiveness shifts corresponded to increases in EC_50_ and decreases in P_max_, respectively. EC_50_ was the half-maximal effective concentration, and P_max_ was the maximum phosphorylation level under each condition. We found that, as a characteristic of FF, the decrease in P_max_ caused by the reduced expression of upstream molecules (IR and IRS2) was propagated downstream as an increase in EC_50_ and a partial recovery of P_max_. In other words, the further downstream the molecule is, the greater the sensitivity shift and the greater the recovery of the responsiveness shift. Furthermore, we also revealed that the magnitude of the responsiveness shift varied depending on the expression levels of downstream molecules. The propagation of the indices varied depending on the parameters and expression levels of the molecules. For example, GSK3β showed an obvious sensitivity shift, while FoxO1 and ACLY exhibited both shifts. In particular, pACLY exhibited a two-step change: a large responsiveness shift immediately after obesity induction, followed by a marked sensitivity shift. Moreover, these characteristics enabled us to reproduce the responses to differences in IR expression levels reported in previous studies. Thus, we believe that our model captures the characteristics of the insulin signaling pathway and the mechanisms of insulin resistance. Furthermore, these analyses revealed that the excessive number of IRs, known as the “spare receptor hypothesis,” is significant because it ensures efficient and robust signal transduction. On the other hand, molecules with an IFFL, such as pS6K and p4E-BP1, showed robust responses despite the attenuation of insulin signaling caused by obesity. We believe that these findings represent one of the mechanisms underlying selective insulin resistance. The insights into the alterations of the insulin signaling pathway due to decreased protein expression levels revealed in this study not only shed light on the mechanisms of insulin resistance, including selective insulin resistance associated with obesity, but also enable a quantitative understanding of the general characteristics observed in signaling pathways.

## 2 Materials and methods

### 2.1 Experimental model and subject details

The institutional animal care and use committee of Kyushu University approved all mouse studies. Male C57BL/6J mice (4 weeks old) were purchased from The Jackson Laboratory Japan. Mice were fed a control diet (Research Diets, Inc., D12450J), and those in the obesity group were switched to a high-fat diet (Research Diets, Inc., D12492) at 5 weeks of age. After overnight fasting, the mice were anesthetized using isoflurane. To suppress endogenous insulin secretion, somatostatin was administered through the jugular vein (3 mg/kg/min). Insulin was administered through the mesenteric vein at the indicated dose (7 ml/kg/min), while maintaining blood glucose levels at a constant level using 20% glucose (CD: 80–150 mg/dL, HFD: 180–250 mg/dL). Blood samples were collected at the indicated time points via the tail vein, and blood insulin levels were measured using a mouse insulin enzyme-linked immunosorbent assay kit (ELISA kit; Fujifilm, 292-89401, 296-89801). At the indicated time points, the mice were sacrificed and their livers were snap-frozen in liquid nitrogen.

### 2.2 Western blotting

Protein extraction and Western blotting were performed as previously described[21]. The following antibodies were used: IR (Cell Signaling Technology, #3025); pIR (Tyr1150/1151, Cell Signaling Technology, #3024); IRS1 (Cell Signaling Technology, #2382); IRS2 (Santa Cruz, sc-8299), Akt (Cell Signaling Technology, #4691); pAkt (Thr308, Cell Signaling Technology, #2965); GSK3β (Cell Signaling Technology, #12456); pGSK3β (Ser9, Cell Signaling Technology, #9323); FoxO1 (Cell Signaling Technology, #2880); pFoxO1 (Ser256, Cell Signaling Technology, #9461); ACLY (Cell Signaling Technology, #4332); pACLY (Ser455, Cell Signaling Technology, #4331); S6K (Cell Signaling Technology, #2708); pS6K (Thr389, Cell Signaling Technology, #9234); 4E-BP1 (Cell Signaling Technology, #9452); and p4E-BP1 (Ser65, Cell Signaling Technology, #9451). To detect the signals of pan- and phospho-specific antibodies on a single membrane, the membrane was incubated in stripping buffer (2% SDS, 62.5 mM Tris pH 6.8, 0.7% 2-mercaptoethanol) for 30 minutes at 50 °C after acquiring the initial luminance data. After confirming that the primary antibodies were removed by treatment with secondary antibodies alone, the membrane was reproved with the other antibody. To compare signal intensities across membranes for the molecules measured, we selected four samples from two time points at each dose from different membranes. These samples were then analyzed together on the same membrane. Simply dividing the luminance signal intensity of the phospho-specific antibody by that of the pan-antibody does not reflect the differences in protein expression across conditions. Moreover, changes in protein expression levels of housekeeping genes have been reported for obesity[22]. Therefore, we normalized the phosphorylation levels using the mean of all data for each condition, as a large dataset was available (CD5w: n_CD5w_ = 81, CD14w: n_CD14w_ = 96, HFD14w: n_HFD14w_ = 96).

The expression ratio of CD14w or HFD14w (Fig. 1G) was defined as follows:

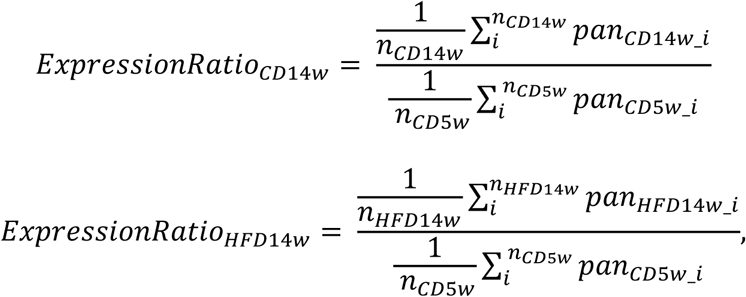

where pan_CD5w_i_, pan_CD14w_i_, and pan_HFD14w_i_ represented the luminance signal intensities of the *i*-th samples of CD5w, HFD14w, and CD14w, respectively. “Pan” refers to signals obtained using pan-specific antibodies. The *i*-th phosphorylated value of the CD5w sample shown in Fig. 1F was calculated as follows:

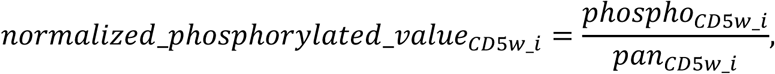

where phospho_CD5w_i_ denoted the luminance signal intensity obtained using phospho-specific antibodies for the *i*-th sample. The *i*-th phosphorylated values of the CD14w and HFD14w samples shown in Fig. 1F were calculated as follows:

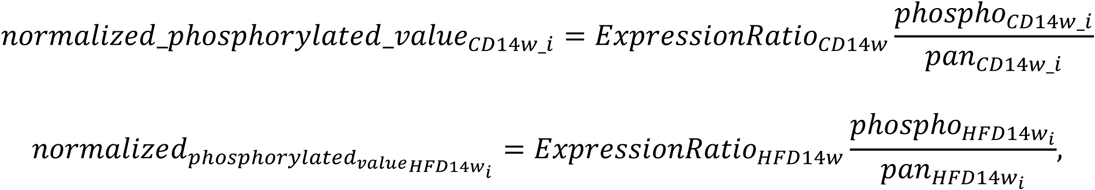

where phospho_CD14w_i_ and phospho_HFD14w_i_ denoted the luminance signal intensities of the *i*-th samples from CD14w and HFD14w, respectively. For 4E-BP1, phosphorylation was evaluated based on the band shifts and the ratio γ/(α+β+γ) was defined as p4E-BP1 in this experiment.

**Fig. 1.**
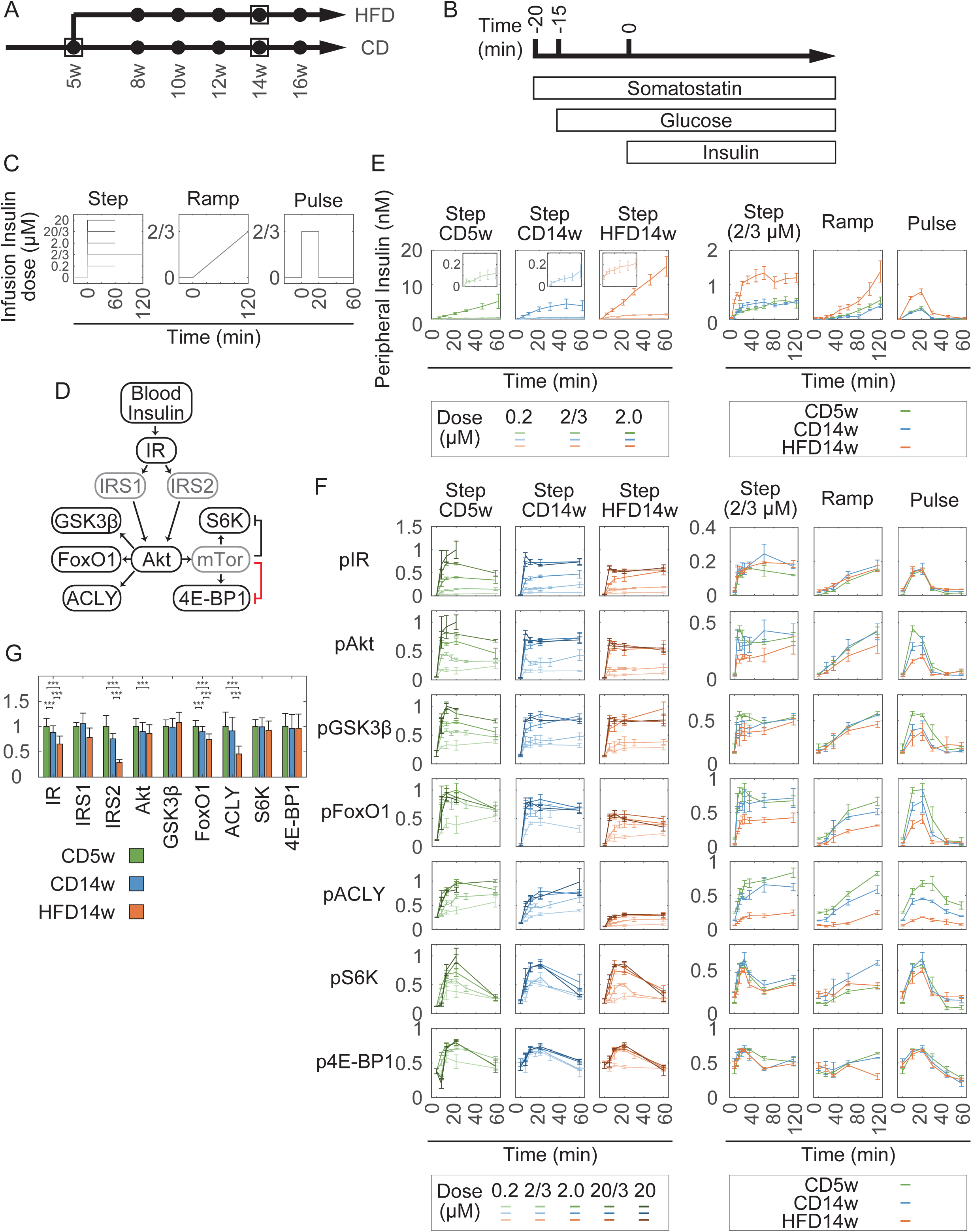
Different temporal patterns of pIR, pAkt, pGSK3β, pFoxO1, pACLY, pS6K, and p4E-BP1 in the liver during obesity progression. (A) Schematic overview of mice used in the intraperitoneal insulin tolerance test (IPITT) and the infusion experiment. Five-week-old mice fed a control diet (CD) and 8, 10, 12, 14, and 16-week-old mice fed either a CD or a high-fat diet (HFD) were used in the IPITT (black circles). For the infusion experiment, 5-week-old mice fed a CD and 14-week-old mice fed either a CD or an HFD were used (white squares). (B) Schematic overview of the procedure for the infusion experiment (see EXPERIMENTAL MODEL AND SUBJECT DETAILS). (C) Insulin infusion patterns. For the step stimulation, 0.2, 2/3, 2.0, 20/3, and 20 μM insulin was injected at a constant rate. The 2/3 μM insulin dose was injected for 120 minutes, and the other doses were injected for 60 minutes. For ramp and pulse stimulations, 2/3 μM insulin was injected according to each pattern. (D) Overview of the insulin signaling pathway. Molecules encircled in black are simulation targets in this study. The red line represents assumed regulation. (E) Time courses of peripheral insulin concentration in the infusion experiments. The time courses of 20/3 and 20 μM step stimulations are shown in Supp. Fig. 2A. Data are shown as mean ± SD. Sample sizes (n) are provided in Star Methods. (F) Time courses of phosphorylation in the insulin signaling pathway during the infusion experiments. Data are shown as mean ± SEM (n = 3). (G) Protein expression levels of each molecule under the indicated conditions. Data are shown as mean ± SD. IR, Akt, GSK3β, FoxO1, ACLY, S6K, and 4E-BP1: n = 114 (CD5w) and n = 117 (CD14w and HFD14w). IRS1 and IRS2: n = 6. Welch’s *t*-test with the Bonferroni method was performed when the differences between the conditions exceeded 15%. *** p < 0.001.

### 2.3 Simulation and parameter estimation

We developed ODE models and performed simulation and parameter estimations using MATLAB (version R2021b, MathWorks). The model structures, reactions, parameters, and differential equations are described in Supp. Fig. 3 and Supp. Tables 1–4. Parameters were estimated as previously described[21]. The insulin concentration administered via the portal vein was calculated based on the injection rate (7 ml/kg/min) and the blood flow rate in the mouse portal vein (72.5 ml/kg/min)[23]. Basal insulin concentration was defined in the model according to previous research[21]. The Mouse Insulin Signaling Model was developed based on our previous insulin-dependent Akt pathway model[21], with four major modifications (Supp. Fig. 3C). First, we modified insulin degradation. Blood insulin levels in the step stimulation showed sustained responses within 60 minutes at 0.2 and 2/3 μM but kept rising at concentrations above 2.0 μM. We attributed this to the limited capacity of insulin-degradation enzymes (IDEs). Therefore, we incorporated an insulin-and-IDE bound state in the new model, the rate-limiting step for insulin degradation (Supp. Fig. 3). Second, we modified the IRS phosphorylation step. Previous models did not distinguish between IRS1 and IRS2. However, we observed that IRS2 expression was more significantly reduced than IRS1 in obese mice. Therefore, we distinguished between the effects of IRS1 and IRS2: pIR phosphorylates both IRS1 and IRS2 and pIRS1 and pIRS2 independently phosphorylate Akt. Third, while our previous model included negative feedback from mTOR to IRS, model comparison based on the BIC showed that the model without negative feedback had a smaller BIC (Supp. Fig. 3B), indicating that it was a better model. Furthermore, some previous studies have questioned the existence of such negative feedback based on transgenic mouse and inhibitor experiments. Therefore, we did not incorporate negative feedback into this model. Finally, we added the Akt downstream molecules, ACLY and 4E-BP1. Phosphorylation of ACLY was modeled as an FF regulation. 4E-BP1 phosphorylation exhibited both a transient (Fig. 1F) and a switch-like response (Fig. 2A). Therefore, 4E-BP1 was incorporated downstream of mTOR as an IFFL regulator with a Hill equation based on the recent findings (Supp. Fig. 3C) regarding its two-step phosphorylation and partial conformational change as an intrinsically disordered protein[24]. The model incorporated phosphorylation of 4E-BP1 at T37/T46 residues, subsequent partial structuring, and phosphorylation at S65/T70 residues (Supp. Fig. 3C). Since the model had a large number of parameters, estimating the optimal parameters at once was challenging. Therefore, as in a previous study[21], the model was divided into modules to reduce the parameter search space, and the final parameter set was estimated.

**Fig. 2.**
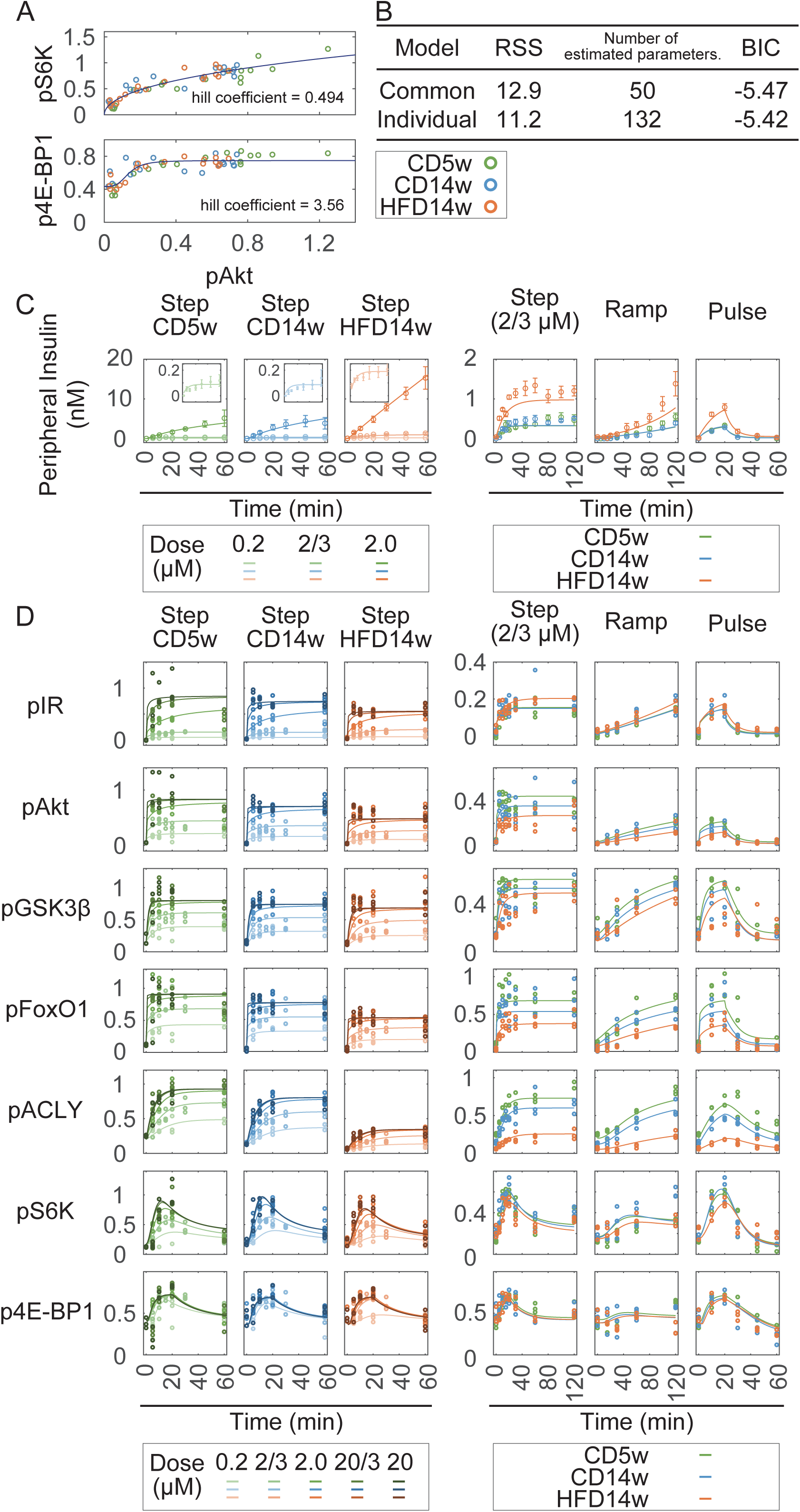
Development of an ordinary differential equation (ODE) model reproducing insulin responses under different conditions. (A) Dose-response curves of pS6K and p4E-BP1 against pAkt. The results of the Hill equation fitting and the experimental data are shown as solid lines and circles, respectively. Data for 0 and 20 minutes were used for the plot. (B) Comparison between the common parameter and individual parameter models. RSS and BIC represent the residual sum of squares and Bayesian information criterion, respectively. For BIC, a smaller value indicates a better model. (C) Time courses of blood insulin levels under the conditions indicated. Solid lines and circles represent simulation and experimental data, respectively. Data are shown as mean ± SD (n=3). (D) Time courses of phosphorylation of the insulin signaling pathway molecules. Solid lines and dots represent simulation and experimental data, respectively.

### 2.4 Toy model analysis

The toy model (Fig. 4A, and Supp. Fig. 5AB) consisted of input, X, and Y, representing a simple signal transduction. The corresponding equations are shown below:

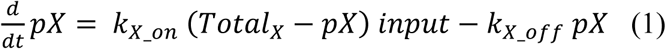

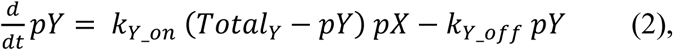

where *Total_X_* and *Total_Y_* were the total expression levels of X and Y; *k_X_on_* and *k_Y_on_* were the phosphorylation rate constants for X and Y; and *k_X_off_* and *k_Y_off_* were the dephosphorylation rate constants for X and Y, respectively.

In the steady state, pX was obtained from Equation (1) as follows:

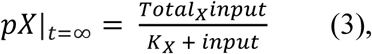

where *K_X_* = *k_X_off_* / *k_X_on_*. In this context, *K_x_* and *Total_X_* represented the EC_50_ and P_max_ of X, respectively. In the steady state, pY was obtained by substituting equation (3) into equation (2) as follows:

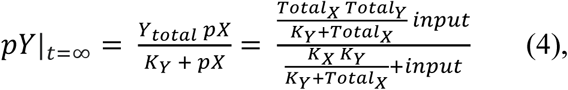

where *K_Y_* = *k_Y_off_* /*k_Y_on_*. The EC_50_ and P_max_ of Y were given as follows:

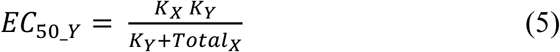

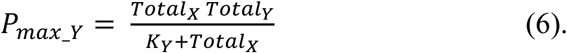

When the expression level of X changed from *Total_X_1_* to *Total_X_2_*, the EC_50_ of X remained unchanged for any *K_X_*, while the P_max_ of X changed from *Total_X_1_* to *Total_X_2_*.

The ratio of the change in the EC_50_ of Y, derived from equation (5), was as follows:

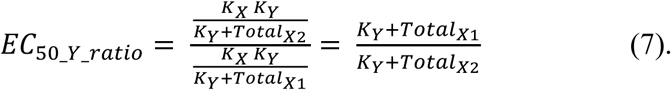

The ratio of the change in the P_max_ of Y, derived from equation (6), was as follows:

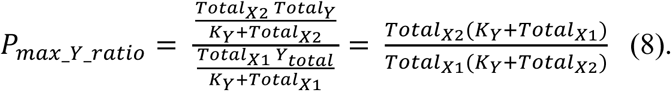

By taking the ratio of P_max_Y_ratio_ to EC_50_Y_ratio_, we obtained the following:

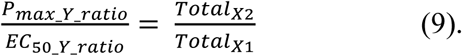

Therefore, P_max_Y_ratio_/EC_50_Y_ratio_ was independent of *K_Y_*.

### 2.5 Fitting experimental data with the Hill equation

For fitting pS6K and 4E-BP1 to pAkt, we focused on the peak of the transient response at 20 minutes under the step stimulation and fitted the responses using the Hill equation as follows:

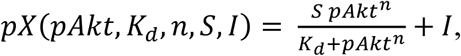

where pX, pAkt, K_d_, n, S, and I represented the phosphorylation levels of the target molecule and Akt, the Hill coefficient, scaling factor, and intercept factor, respectively.

For fitting pIR, pAkt, pGSK3β, pFoxO1, and pACLY to blood insulin levels, we considered the phosphorylation levels of each molecule at 0 minutes, 0.2, 2, 20/3, and 20 μM at 60 minutes, and 2/3 μM at 120 minutes under the step stimulation as the steady state and then used these data for fitting. To ensure comparability with the constructed model, which did not incorporate cooperative regulation, the data were fitted using the Hill equation, with the Hill coefficient fixed at 1, as follows:

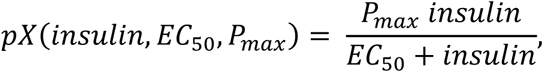

where pX and insulin represented the phosphorylation levels of the target molecules and peripheral insulin levels (nM), respectively. The EC_50_ and P_max_ values were determined separately for each feeding condition and for each molecule.

### 2.6 Calculation of insulin demand

First, we determined the steady-state phosphorylation levels of the target molecules at 0.1 nM insulin in 5-week-old mice. Next, we simulated the blood insulin levels required for the target molecule to reach the same phosphorylation level for each feeding condition and week of age.

## 3 Results

### 3.1 Age-dependent attenuation of insulin signaling in the liver during obesity progression induced by a high-fat diet

We examined the progression of insulin resistance in mice after obesity was induced by an HFD. After one week of acclimation, mice were randomly divided into either a control diet (CD) group or an HFD group at 5 weeks of age. Then, the mice were analyzed at 5, 8, 10, 12, 14, and 16 weeks of age. First, we examined physiological changes in CD and HFD mice under overnight fasting conditions. Compared to CD mice, HFD mice showed marked increases in body weight, fasting blood glucose, and fasting blood insulin levels (Supp. Fig. 1A–C). Notably, an evaluation of insulin resistance progression showed that these values increased rapidly in the HFD group between 12 and 14 weeks of age. Next, mice were injected intraperitoneally with 0.7 U/kg insulin for the intraperitoneal insulin tolerance test (IPITT), and blood glucose and insulin levels were measured for 20 minutes (Supp. Fig. 1D–F). In HFD mice, fasting blood insulin levels increased with week of age, while the hypoglycemic effect was suppressed, indicating the insulin resistance progression associated with obesity. The expression and phosphorylation levels of the IR and Akt were examined using Western blotting (Supp. Fig. 1G–K) with livers obtained at 20 minutes. No significant difference in phosphorylation was observed between CD and HFD mice. The efficiency of signal transduction can be thought of as how much an activated upstream molecule can activate a downstream molecule. Therefore, to represent signal transduction efficiency, we calculated the ratio of the amount of phosphorylation of upstream molecules to that of downstream molecules. Signal transduction efficiency significantly decreased from IR to Akt from 10 weeks of age in HFD mice, indicating attenuation of insulin signaling from IR to Akt (Supp. Fig. 1L).

### 3.2 Alterations of the insulin signaling pathway in the liver of mice during obesity progression

In our previous studies, we showed that the insulin signaling pathway processes blood insulin patterns and selectively regulates downstream molecules depending on the temporal patterns of insulin[21,25]. Therefore, in this study, we examined how this selective regulation is altered during obesity progression. To examine the response of insulin signaling in vivo, we administered insulin via the mesenteric vein or portal vein (infusion experiment) to mimic physiological insulin secretion, as previously described[21]. Based on the results of the IPITT, which showed that fasting blood glucose and fasting blood insulin levels in HFD mice increased significantly between 12 and 14 weeks of age (Supp. Fig. 1A–C), we used 14-week-old CD and HFD mice and 5-week-old CD mice (Fig. 1A square) for the infusion experiment. Somatostatin was administered intravenously from 20 minutes before insulin stimulation to suppress endogenous insulin secretion. To maintain blood glucose levels, glucose was also administered intravenously from 15 minutes before insulin infusion (Fig. 1B). Insulin was administered in three different patterns (step, ramp, and pulse). Five concentrations, ranging from 0.2 to 20 µM, were used for the step stimulation (Fig 1C). In addition to IR, Akt, GSK3β, FoxO1, and S6K, which were examined in the previous study[21], we also measured ACLY and 4E-BP1, both of which are associated with obesity (Fig 1D–F, and Supp. Fig. 2).

Peripheral insulin levels in the step stimulation showed a sustained response within 60 minutes at 0.2 and 2/3 μM but tended to increase continuously at concentrations higher than 2.0 μM (Fig. 1E and Supp. Fig. 2A). Since 20 μM stimulation in CD5w led to the death of some mice between 20 and 60 minutes, only the data within 20 minutes were analyzed in subsequent analyses. These results indicate that, at least in CD5w mice, 20 μM insulin stimulation is not physiological. In HFD14w mice, compared to CD14w mice, expression levels of the proteins IR, IRS2, FoxO1, and ACLY were significantly decreased by 25%, 62%, 17%, and 50%, respectively (Fig. 1G). The decreased expression of IR, IRS2, and ACLY in obesity is consistent with previous results[8,10,26]. Given that FoxO1 expression is regulated via the cAMP-PKA pathway[27], and that this pathway is downregulated in obese mice[28], this observed decrease in FoxO1 expression may be attributable to cAMP-PKA pathway suppression. On the other hand, the qualitative temporal patterns of these molecules were similar between mouse conditions. In particular, pIR, pAkt, pGSK3β, and pFoxO1 showed sustained responses, while pS6K showed a transient pattern, consistent with our previous results[21]. These findings indicate that despite obesity-induced insulin resistance, the qualitative characteristics are conserved. In the case of quantitative changes, although peripheral insulin levels were higher in HFD14w mice than in CD14w mice under 2/3 µM insulin stimulation, phosphorylated IR was similar between the two groups (Fig. 1EF). However, despite the same level of IR phosphorylation, Akt phosphorylation was reduced. The Akt downstream molecules, GSK3β, FoxO1, and ACLY, showed a decrease in phosphorylation similar to that of Akt, but the extent of the attenuation differed, with a particularly strong decrease in ALCY. S6K and 4E-BP1 showed similarly transient activation in both groups. Peak phosphorylation of S6K increased in a concentration-dependent manner in response to step stimuli ranging from 0.2 to 2.0 μM, whereas 4E-BP1 exhibited a switch-like response. Even under 0.2 μM step stimulation, all CD14w mice exhibited transient responses, while none of the HFD14w mice did. Some CD5w mice showed transient responses, whereas others did not (Supp. Fig. 2B). These results indicate that 4E-BP1 responded to insulin in a switch-like manner independent of mouse conditions and insulin concentration, while other molecules responded in a concentration-dependent manner.

### 3.3 Development and evaluation of an ordinary differential equation model that recapitulates alterations in the insulin signaling pathway in CD and HFD mice

We developed an ordinary differential equation (ODE) model that reproduced the dynamics of the insulin signaling molecules under each condition by introducing four modifications to our previous model[21] (Supp. Fig. 3). (1) Sustained increases in peripheral insulin levels were observed in the step stimulation with high insulin concentrations. We considered this to be due to the limited degradation capacity of insulin-degrading enzymes. Therefore, we incorporated an insulin and insulin-degrading enzyme bound state (enzyme–substrate complex) into the model. This provided an upper limit to insulin degradation determined by the turnover rate of the insulin-degrading enzyme. (2) The decrease in IRS1 and IRS2 protein levels due to obesity differed (Fig. 1G). Therefore, we distinguished between the effects of IRS1 and those of IRS2 in the model. (3) We examined the contribution of feedback from mTOR to IRS using mathematical models and found no significant difference (Supp. Fig. 3A); therefore, we did not consider this feedback in the model. (4) ACLY and 4E-BP1, which were not included in the previous study[21], were incorporated into the model. Since 4E-BP1 showed a transient response similar to that of S6K, we assumed IFFL regulation as with S6K (Fig. 1D, red line). However, unlike S6K, the 4E-BP1 band shift showed a switch-like response (Fig. 1F and Supp. Fig. 2B). Therefore, we focused on the peak of the transient response at 20 minutes after the step stimulation and fitted the response to phosphorylation of Akt, an upstream molecule, using the Hill equation (Fig 2A). As a result, the Hill coefficient of S6K was less than 1, indicating a non-cooperative response, while that of 4E-BP1 was 3.6, suggesting cooperative phosphorylation regulation. In other words, this cooperative phosphorylation resulted in the same transient 4E-BP1 response once Akt phosphorylation exceeded a certain threshold. This result was also confirmed when the phosphorylation of S65 on 4E-BP1 was measured instead of the band shift (Supp. Fig. 2D). The two-step phosphorylation reaction has previously been reported for 4E-BP1, an intrinsically disordered protein, in which T37 and T46 induces partial structuring, followed by phosphorylation at S65 and T70[24]. Therefore, we assumed that this two-step phosphorylation accounted for the cooperativity and incorporated both IFFL and Hill regulation into the model for 4E-BP1.

Decreased protein expression and altered dephosphorylation activity may contribute to insulin resistance[29,30]. To quantitatively analyze the mechanisms underlying obesity-induced insulin resistance, we first examined the impact of changes in protein expression levels and other contributions using the ODE model. We developed two models: an individual-parameter model, which allowed both protein expression levels and rate constants to vary under each condition; and a common-parameter model, in which only protein expression levels differed (Supp. Table 3). We compared the two models using the Bayesian information criterion (BIC) and found that the common-parameter model provided a better fit (Fig. 2B). These results suggest that changes in protein levels were the primary cause of obesity-induced insulin resistance, at least up to 9 weeks after obesity induction. As shown in Fig. 2CD, and Supp. Fig. 2A, the simulations using the common-parameter model successfully reproduced the experimental results. Therefore, the following analyses were performed using the common-parameter model named “Mouse Insulin Signaling Model.”

### 3.4 Quantitative comparison of obesity-induced attenuation of insulin signaling

To compare the effects of obesity on each molecule in the insulin signaling pathway, we focused on the response to 2/3 uM step stimulation at 20 minutes and compared the phosphorylation ratios of each molecule between CD14w and HFD14w mice (Fig. 3A). A gradual decrease in the efficiency of signaling from blood insulin to IR and Akt was observed in obese mice in both the simulation and the experiment, despite increased peripheral insulin levels. However, the phosphorylation ratios of Akt downstream molecules varied: the phosphorylation ratios of pGSK3β, pS6K, and p4E-BP1 tended to be larger than that of pAkt. These results indicate that signaling efficiency up to Akt, which was impaired by obesity, was partially restored in downstream molecules. Conversely, the phosphorylation of ACLY was markedly attenuated to approximately half that of the control, indicating that the impact of obesity on insulin signaling differed depending on the molecule (or pathway).

**Fig. 3.**
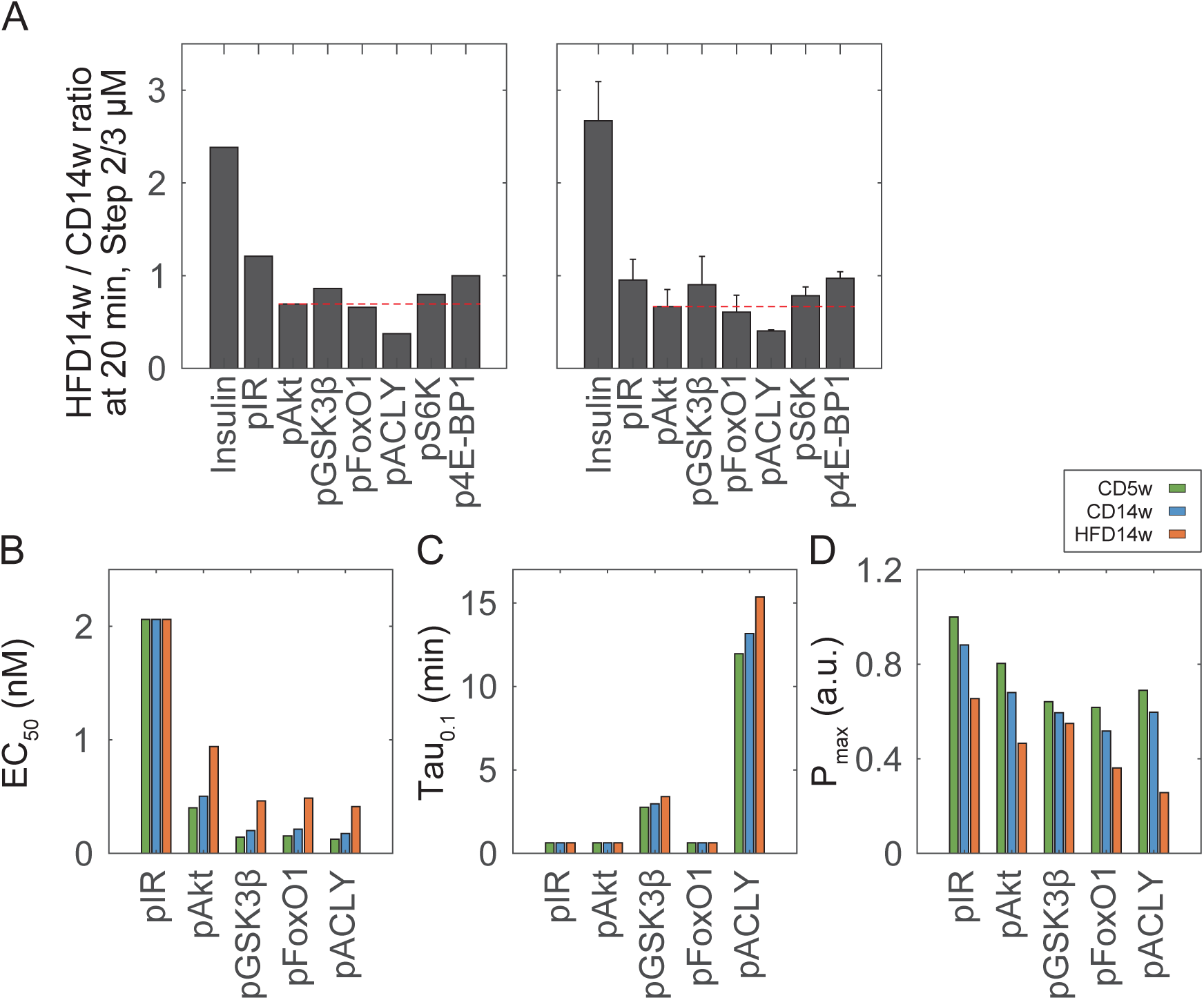
Obesity-induced information attenuation of insulin signaling. (A) HFD14w to CD14w phosphorylation ratios of molecules in the insulin signaling pathway at 20 minutes in the 2/3 μM insulin step stimulation: simulation (left) and experiment (right). Data are shown as mean ± SD (n=3). (B) EC_50_; (C) Tau_0.1_; and (D) P_max_ for each molecule were calculated from the simulation. Tau_0.1_ represents the time constant at 0.1 nM blood insulin concentration. P_max_ represents the theoretical maximum phosphorylation of each protein, with the maximum value under the CD5w condition set to 1.

In a previous study, we found that each molecule in the insulin signaling pathway can selectively regulate downstream molecules by processing information encoded in the concentration and temporal changes (increasing rate) of blood insulin[21,25]. In this study, we also focused on the EC_50_ and tau, which are indices of concentration and response speed in FF regulation, to examine how these indices are altered in obese mice with insulin resistance (Fig. 3BC). First, we examined EC_50_ (Fig. 3B), which represents the half-maximal effective concentration. Under CD5w, CD14w, and HFD14w conditions, the EC_50_ values of IR were the same. On the other hand, those of Akt and its downstream molecules were higher in the HFD14w group than in the CD5w and CD14w groups. Interestingly, the EC_50_ of IR (2.06 nM) was higher than the postprandial blood insulin level in healthy mice[31,32]. This implies that more than half of the IRs were not phosphorylated even after insulin secretion after feeding. Next, we examined the time constant (tau), which indicates how rapidly downstream molecules can follow the changes in upstream molecules: molecules with a small tau can rapidly follow an upstream molecule. Since tau varies depending on stimulus concentration, we defined Tau_0.1_ as the time required to reach 63.2% of the steady state in response to a blood insulin concentration of 0.1 nM (Fig. 3C) and used it for comparison. The Tau_0.1_ of pIR, pAkt, pGSK3β, and pFoxO1 was small, whereas that of pACLY was larger than that of other Akt downstream molecules. Similar results were observed for Tau_1_, given a blood insulin concentration of 1 nM (Supp. Fig. 4). These results indicate that pACLY could not respond to rapid changes in insulin concentration and received less information about the increasing rate than other molecules.

Furthermore, to compare the phosphorylation of the same molecules under different conditions, we defined P_max_ as the phosphorylation level when the blood insulin level was assumed to be infinite. P_max_ was normalized so that complete phosphorylation of the total expression level in CD5w was set to 1 (Fig. 3D). Therefore, P_max_ allowed for direct comparison of the expression and phosphorylation levels of the same molecule under different conditions. Changes in insulin signaling under steady-state conditions were examined using the two indices, P_max_ and EC_50_. The P_max_ of all molecules was lower in HFD mice than in CD5w and CD14w mice (Fig. 3D), although the decrease was less pronounced for GSK3β. In other words, the maximum possible phosphorylation (dynamic range) was reduced for many molecules in HFD mice.

### 3.5 Investigation of the EC_50_ and P_max_ transmission characteristics in the signaling pathway

Our finding that the common parameter model can reproduce the responses of each signaling molecule indicates that differences in protein expression levels account for differences in signaling under both CD and HFD conditions. Therefore, we investigated the characteristics of FF, which constitute a substantial part of the regulation of the insulin signaling pathway examined in this study, by analyzing how changes in upstream protein expression influenced the EC_50_ and P_max_ of downstream molecules. The attenuation of the insulin effect on hypoglycemia can be characterized by two aspects, sensitivity shift and responsiveness shift[2,19,20], which correspond to an increase in the EC_50_ and a decrease in P_max_, respectively (Fig. 4A). We also applied these concepts to the analysis of molecules with FF. Since the Mouse Insulin Signaling Model describes FF using mass action kinetics without enzyme–substrate complexes, the EC_50_ can be derived analytically (see Star Methods). Therefore, before investigating the characteristics of the insulin signaling pathway, we examined the characteristics of a signaling pathway with FF using a toy model consisting of three components, Input, X, and Y. In this toy model, the steady-state values of X and Y were given by the following equations (Fig. 4A, see Star Methods):

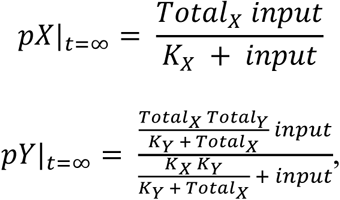

where *K_X_* = *k_X_off_*/*k_X_on_*, and *K_Y_* = *k_Y_off_*/*k_Y_on_*, *K^X^* and *Total_X_* (the total amount of X) corresponded to the EC_50_ and P_max_ of X, respectively; similarly, *K_X_K_Y_*/(*K_Y_* + *Total_X_*) and *Total_X_Total_Y_*/(*K_Y_* + *Total_X_*) corresponded to the EC_50_ and P_max_ of Y, respectively. Therefore, the EC_50_ of X remained unchanged regardless of changes in its expression level, whereas a decrease in X expression led to a reduction in its P_max_. This is consistent with the result that the EC_50_ of IR remained unchanged for each condition under different IR expression levels (Fig. 3B). In contrast, since both the EC_50_ and P_max_ of the downstream molecule Y depended on *Total_X_*, changes in the upstream molecule propagated and influenced both the EC_50_ and P_max_ of Y. For example, consider the case where the expression level of upstream molecule X decreased from *Total_X1_* to *Total_X2_*. For molecule X, the ratio P_max_X2_/P_max_X1_ (P_max_X_ratio_) decreased, whereas the ratio EC_50_X2_/EC_50_X1_ (EC_50_X_ratio_) remained the same (Supp. Fig. 5A). In contrast, for molecule Y, regardless of *K_Y_*, the EC_50_ ratio (EC_50_Y2_/EC_50_Y1_; EC_50_Y_ratio_) increased and the P_max_ ratio (P_max_Y2_/P_max_Y1_; P_max_Y_ratio_) became greater than *Total_X2_*/*Total_X1_* (Supp. Fig. 5B). Since P_max_X_ was equal to *Total_X_*, P_max_X2_/P_max_X1_ (P_max_X_ratio_) was equal to *Total_X2_*/*Total_X1_*. Thus, P_max_Y_ratio_ was always greater than P_max_X_ratio_, indicating that P_max_ always recovered in the downstream molecule. Additionally, the ratio of P_max_Y_ratio_ to EC_50_Y_ratio_ became independent of *K_Y_* and equaled *Total_X2_*/*Total_X1_* (Supp. Fig. 5B, right and Star Methods). In other words, there was a trade-off relationship between the alterations in EC_50_Y_ratio_ and P_max_Y_ratio_ of downstream molecules, which was dependent on the change in X expression levels.

In the Mouse Insulin Signaling Model, increases in EC_50_ and decreases in P_max_ were observed for Akt and its downstream molecules under HFD conditions (Fig. 3BC). To validate these findings, we examined whether similar trends could be observed directly in the experimental data (“static-fitting”) without relying on the ODE model (“ODE-fitting”) (Fig. 4B; see Star Methods). As a result, the EC_50_ and P_max_ values obtained by static-fitting also showed an increase and a decrease, respectively, and were comparable to those obtained by ODE-fitting (Fig 4C). These results indicate that the signaling pathway characteristics identified using the toy model were also evident in the actual insulin signaling pathway. Next, we examined how the EC_50_ and P_max_ of the downstream molecules were altered depending on the changes in IR and Akt expression levels (Supp. Fig. 5C and Fig. 4D). As IR or Akt expression levels decreased, the increase in EC_50_ was accelerated in downstream molecules and the recovery of P_max_ was greatly reduced. In other words, the higher the IR expression level, the more robust the characteristics (EC_50_ and P_max_) of downstream molecules became. For both EC_50_ and P_max_, the slope of the original expression level (at a fold change of 1) was steeper for Akt than for other molecules when IR expression was changed. Furthermore, the responses of pGSK3β, pFoxO1, and pACLY to changes in Akt expression were also steeper than their responses to changes in IR expression. These findings indicate that alterations in upstream molecules weakened/recovered as they propagated to downstream molecules.

**Fig. 4.**
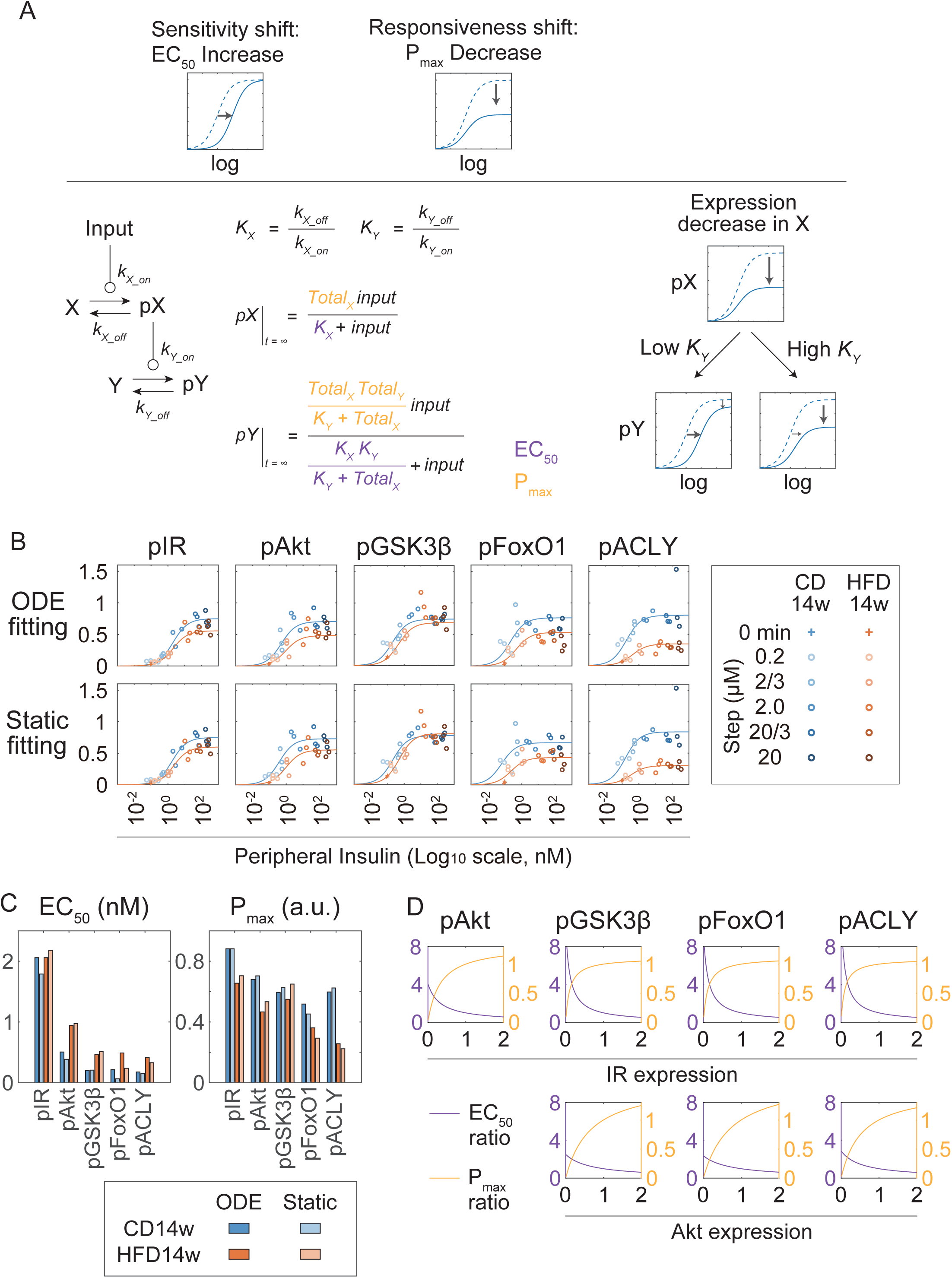
Transmission characteristics of the EC_50_ and P_max_ of downstream molecules in the insulin signaling pathway. (A) Characteristics of the sensitivity shift (increased EC_50_) and responsiveness shift (decreased Pmax) in signaling pathways revealed by the toy model analysis. A reduction in the expression of upstream molecules induces a sensitivity shift, while the decrease in P_max_ of the upstream molecules due to reduced expression is partially compensated in the downstream molecules, depending on the value of *K_Y_*. (B) Dose–response curve of each molecule against blood insulin levels, as estimated by the Mouse Insulin Signaling Model (ODE-fitting) or calculated directly from experimental data (static-fitting). Dots indicate experimental data for 0 minutes, 60 minutes (0.2, 2.0, 20/3, and 20 μM), and 120 minutes (2/3 μM). (C) EC_50_ and P_max_ values obtained from the Mouse Insulin Signaling Model (ODE-fitting) or directly from experimental data (static-fitting). (D) The relationship between the expression levels of IR and the EC_50_ or P_max_ of the downstream molecules simulated by our model. Based on the estimated parameters, the IR-expression-dependent EC_50_ and P_max_ of each molecule are shown. The EC_50_ and P_max_ for the IR expression of CD14w mice were normalized to 1. The EC_50_ and P_max_ values are indicated in dark blue and yellow, respectively.

We also analyzed how changes in the expression levels of upstream molecules affected the tau of downstream molecules (Supp. Fig. 5D). When IR expression levels were varied, the effects differed depending on the tau of each downstream molecule: molecules with a larger tau (e.g., pACLY) were more affected. On the other hand, for all molecules, a decrease in the expression levels of upstream molecules led to an increase in tau. As the expression levels of upstream molecules increased, changes in tau became smaller, indicating that the higher the IR expression level, the more robust the tau of the downstream molecules became.

### 3.6 Investigation of changes in the EC_50_ and P_max_ during obesity progression

The ability of the common model to reproduce the observed responses, along with the toy model analysis, indicated that, given the expression levels of each molecule, the EC_50_ and P_max_ could be calculated using the parameters of the Mouse Insulin Signaling Model. Therefore, we calculated the EC_50_ and P_max_ of each molecule at each week of age during obesity progression, based on the expression level data obtained from mice used in the IPITT experiment (Supp. Fig. 6AB), and examined how these values changed over time (Fig. 5A). In obese mice, the EC_50_ of pAkt, pGSK3β, pFoxO1, and pACLY increased in a week-age-dependent manner, especially between 12 and 14 weeks of age, showing nearly a 2-fold increase. On the other hand, the change in the P_max_ was similar to the change in the expression levels of the respective molecules (Fig. 5A and Supp. Fig. 6B). The P_max_ was also dependent on its expression levels. Therefore, to isolate the effect of decreases in the upstream molecules, we calculated the changes in the P_max_, excluding the contribution of each molecule’s own expression level (Fig. 5A, lower panel; Star Methods). We found that, compared to pAkt, the changes in P_max_ of pGSK3β, pFoxO1, and pACLY in response to alterations in the expression levels of upstream molecules were smaller. This indicates that the effect of upstream molecule reduction on the P_max_ was weaker in molecules located further downstream. These findings indicate that changes in the P_max_ of Akt’s downstream molecules were primarily dependent on changes in their own expression levels and were less influenced by changes in IR and IRS2 expression levels. In particular, the P_max_ of pACLY was already reduced to its minimum level by eight weeks of age, immediately after the switch to an HFD, and this reduction was largely attributable to the decrease in ACLY expression itself (Supp. Fig. 6B). While the P_max_ of downstream molecules recovered, the EC_50_ of pAkt, pGSK3β, pFoxO1, and pACLY largely increased in HFD mice, which was caused by the decreases in the expression levels of IR and IRS2 (Supp. Fig. 6B). In addition, fasting blood insulin levels increased in the same week (Supp. Fig. 1C), and negative correlations were observed between fasting blood insulin levels and expression levels of IR and IRS2 (Fig. 5B). These findings suggest that hyperinsulinemia might have served as a compensatory response to the increase in the EC_50_ caused by reductions in IR and IRS2 expressions. To examine the effect of increased blood insulin levels on signaling compensation, we determined the phosphorylation levels of each molecule at blood insulin levels of 0.1 nM in CD5w and calculated the insulin concentration required to achieve the same phosphorylation levels under each condition (“insulin demand”) (Fig. 5C). The results showed that the insulin demands of IR, Akt, GSK3β, and FoxO1 increased between 12 and 14 weeks of age in HFD mice. Insulin demands were correlated with fasting insulin concentrations in HFD mice (Fig. 5D), suggesting that the increase in the EC_50_ associated with the decrease in upstream molecules might have led to an increase in the optimal insulin concentration and, therefore, an increase in blood insulin levels. On the other hand, ACLY could no longer be compensated from eight weeks of age (Fig. 5C). Among IR, IRS, and ACLY, which decreased markedly in HFD mice (Fig. 1G), only ACLY did not show a significant correlation between its expression level and fasting blood insulin concentration (Fig. 5B). These results indicate that the responsiveness shift of pACLY was independent of changes compensated for by hyperinsulinemia.

**Fig. 5.**
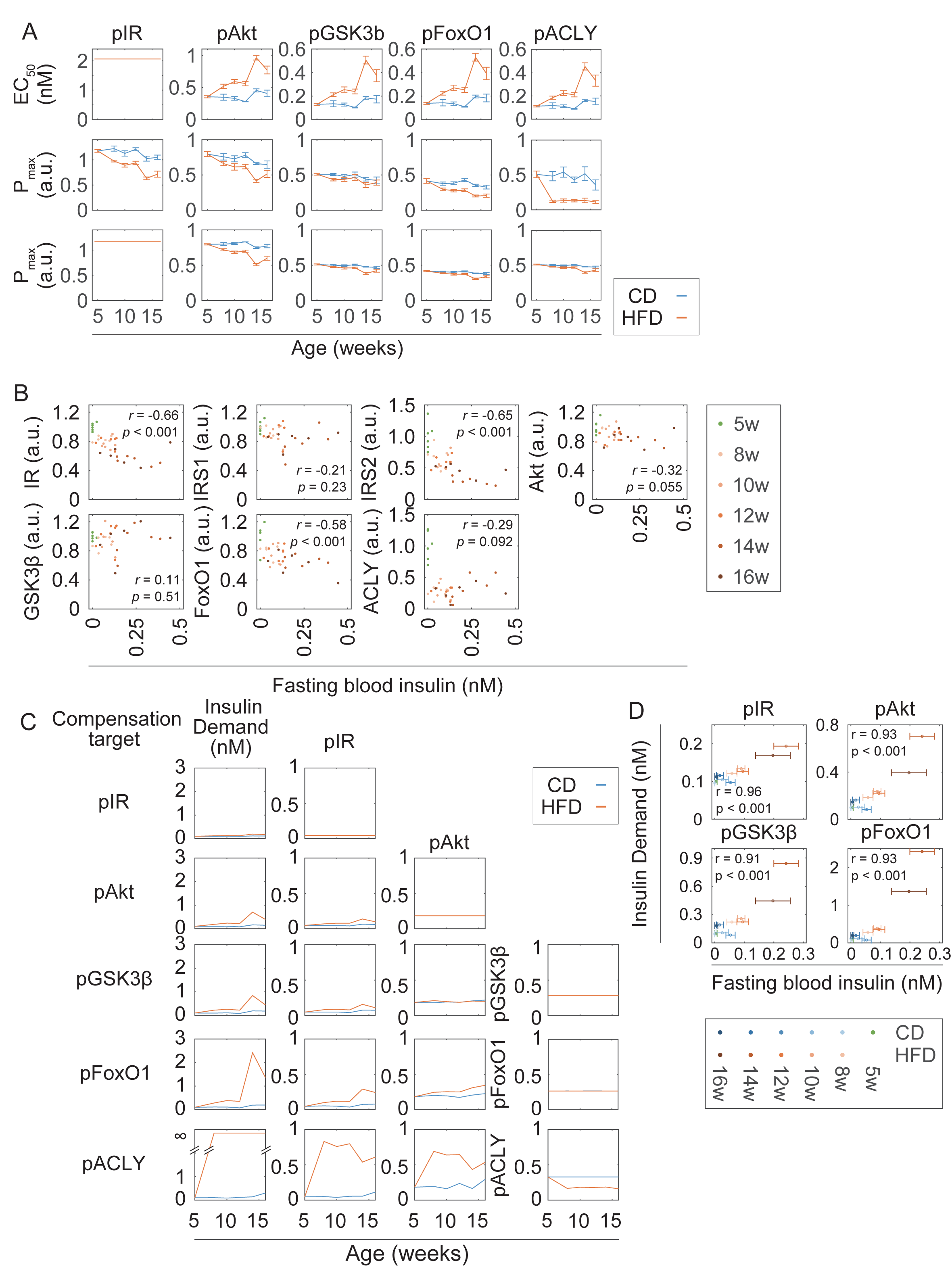
Week-of-age-dependent attenuation of insulin signaling during obesity progression. (A) Alterations in the EC_50_ and P_max_ values of each molecule during obesity progression. EC_50_ (upper panel) and Pmax (middle panel) values were calculated from the protein expression data at the indicated time points with the estimated parameters. The lower panel shows the P_max_ values calculated without the contribution of changes in their protein expression levels. Data are shown as mean ± SD (n=6). (B) Scatter plots of individual protein expression levels against fasting blood insulin levels. (C) Calculated insulin demand (left) required to compensate for the response of the indicated molecule and the corresponding response of each molecule. See Star Methods for details (“Calculation of insulin demand”). If compensation was not possible, insulin demand was set to infinity. (D) The relationship between calculated insulin demand and fasting blood insulin levels under each mouse condition.

### 3.7 Mouse Insulin Signaling Model reproduced previously reported downstream responses to perturbation and mutations in IR

The contribution of IR to the regulation of downstream molecules and responses has been investigated[3,4] using perturbations and mutations in IR. Our analysis using the Mouse Insulin Signaling Model enabled us to quantitatively investigate how the effects of IR propagate to downstream molecules by mathematically evaluating the EC_50_ and P_max_. Therefore, we examined the insulin responses in previous reports.

Kono et al. reported that a 2.4% reduction in IR expression by trypsin treatment increased the EC_50_ for glucose oxidation by 20- to 30-fold, without causing a responsiveness shift[12]. Our toy model analysis showed that the P_max_ratio_/EC_50_ratio_ equaled *Total_X2_*/*Total_X1_*, regardless of the parameters (Supp. Fig. 5B). This implies that even when IR expression was reduced to 2.4% (P_max_ratio_), the downstream P_max_ could be maintained by increasing the EC_50_ by approximately 40-fold (EC_50_ratio_), regardless of how many steps downstream the response occurred. This increase in EC_50_ (the sensitivity shift) is generally consistent with the value reported by Kono et al. Note that this compensation is attributed to the coordinated response of the signaling pathway, rather than an excessive amount of IR expression alone. Merry et al. reported that mice heterozygous for IR showed no abnormalities in Akt phosphorylation and maintained normal glucose and insulin tolerance[13]. In these IR+/- mice, IR expression levels were reduced by approximately half, while IRS2 RNA levels were approximately doubled. To simulate the condition, we used the Mouse Insulin Signaling Model with 0.5-fold IR and two-fold IRS2 and found that both the EC_50_ and P_max_ of Akt phosphorylation were almost restored (Supp. Fig. 6C). This may explain the absence of glucose and insulin intolerance. Several mutations that impair IR activity and are associated with abnormal glucose metabolism have been identified in humans[3,4]. To investigate this using the Mouse Insulin Signaling Model, we simulated a 0.5-fold reduction in IR activity (*k_on_*) and examined the downstream response (Supp. Fig. 6D; see also Star Methods). We found that the EC_50_ of all downstream molecules, including IR, were doubled without affecting their P_max_. In other words, unlike a decrease in upstream molecule expression, a decrease in enzyme activity was entirely propagated to downstream molecules, resulting in increased EC_50_. Indeed, patients with mutations in IR activity exhibit insulin resistance[3,4] but this may be a different mechanism from the decrease in the IR or IRS2 expression due to obesity.

### 3.8 A large EC_50_ of IR ensured robust signal transduction

The EC_50_ of IR under each condition, calculated using the ODE model and static-fitting, was as follows: 2.06 nM (ODE-fitting: CD14w and HFD14w) (Fig. 3B); 1.79 nM (static-fitting: CD14w); and 2.18 nM (static-fitting: HFD14w) (Fig. 4C). These values are higher than the blood insulin levels in normal mice during feeding. This indicates that more than half of the IR remains unphosphorylated even when insulin is secreted during feeding. This phenomenon, the existence of an excessive number of receptors, is known as the “spare receptor hypothesis,” and has been reported for many receptors[14,15]. The maintenance of a large pool of unphosphorylated receptors incurs a biological cost. Why do cells bear this cost? To address this, we investigated the potential advantages of this property.

In the Mouse Insulin Signaling Model, EC_50_ against the upstream molecule can be expressed as the ratio of the dephosphorylation to phosphorylation rate constants (*k_off_*/*k_on_*) (Fig. 4A). Therefore, we varied the *k_off_* of IR (Supp. Table 3, *k7*) and examined how changes in EC_50_ influenced downstream molecules. Although the changing of *k_off_* affects both EC_50_ and tau, we focused solely on steady-state responses in this analysis; thus, we ignored the influences on tau. When the EC_50_ of IR was lowered by half, the EC_50_ of the downstream molecules also decreased (Fig. 6A). As shown in the toy model analysis (Fig. 4A), even if the EC_50_ of X (EC_50_X_) decreased, it was possible to maintain the EC_50_ of Y (EC_50_Y_) by increasing the *K* of Y (*KY*). Therefore, we searched for a condition under which the EC_50_ of Akt equaled the original parameter set in CD14w by increasing the *k_off_* of Akt (Supp. Table 3, *k14*), while keeping the EC_50_ of IR reduced to half of the original parameter set (Fig. 6B). Under this condition, the P_max_ of Akt decreased (Fig. 6C). This corresponds to the finding of the toy model analysis that increasing *K_Y_* to raise the EC_50_ of Y weakened the recovery of P_max_ of Y. This result indicates that even when the expression level of Akt remained unchanged and the input sensitivity (EC_50_) was the same, the output dynamic range (P_max_) became narrower, indicating a decrease in signaling capacity. Even under this condition, increasing Akt expression can restore the Akt P_max_ to its original value; however, this comes at another biological cost. To further investigate the effect of reduced IR expression in obesity, we analyzed how decreasing IR expression levels affected downstream molecules under the conditions described above (IR EC_50_ halved and Akt EC_50_ maintained) (Fig. 6D). Under this condition, sensitivity to changes in IR expression was higher than that under the original condition: the P_max_ of downstream molecules decreased more and the EC_50_ increased less when IR expression was reduced. This result corresponds to that shown in Fig. 6B: even if the EC_50_ of Akt under CD conditions was restored by changing the *k_off_* of Akt, the HFD group was not fully restored to its original value. Furthermore, we performed a signal compensation analysis as shown in Fig. 5C. Under the changed condition (solid line), a higher insulin level was required for compensation, compared to the original condition (Fig. 6E). These results indicate that, when the EC_50_ of IR was small, a decrease in IR expression necessitated costly compensation to restore downstream P_max_.

**Fig. 6.**
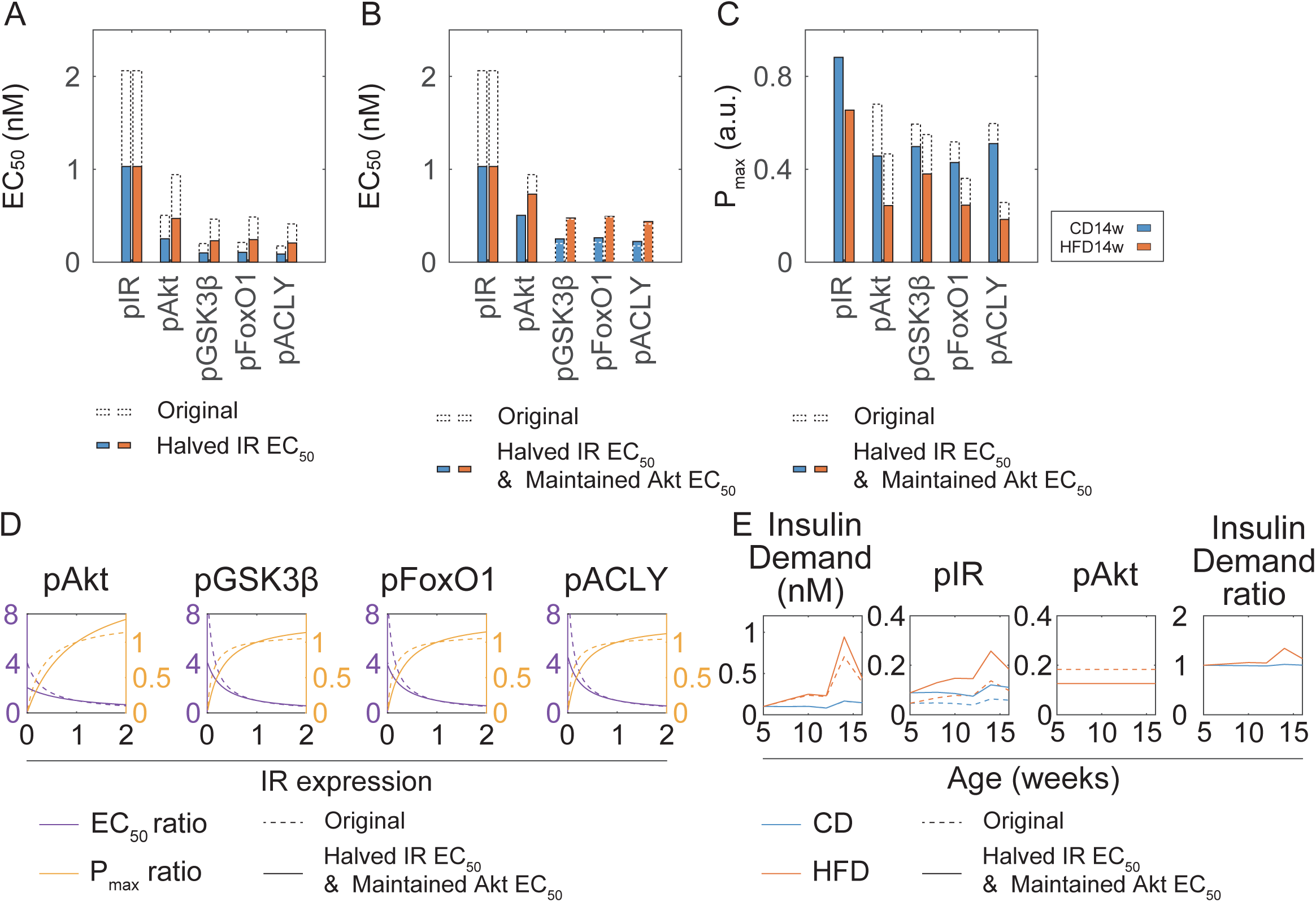
Spare insulin receptors ensure robust signal transduction. (A) Effects on downstream molecules when the EC_50_ of IR is halved. Dotted and solid lines indicate the original EC_50_ and the halved EC_50_, respectively. (B) EC_50_; and (C) P_max_ values when the EC50 of Akt under CD14w is restored to its original value under the above condition. Dotted and solid lines indicate the original EC_50_ and the halved EC_50_, respectively. (C) IR-expression-dependent changes in the EC_50_ and P_max_ of downstream molecules when the EC_50_ of IR is halved. The EC_50_ and P_max_ values are indicated in dark blue and yellow, respectively. Under both conditions, the EC_50_ and P_max_ values of IR expression under the CD14w condition were normalized to 1. Dotted and solid lines indicate the original and changed conditions, respectively. (D) Comparison of insulin demand when the EC_50_ of IR is halved. See Star Methods for details (“Calculation of insulin demand”). The ratio of insulin demand under the original condition to that under changed conditions is shown on the right.

### 3.9 IFFL regulation enabled S6K and 4E-BP1 to be robust to the changes caused by obesity

Insulin secretion exhibits several patterns, including a 10- to 15-minute pulsatile secretion, additional secretion (2-hour period), and basal secretion (a daily period)[33–36]. Therefore, in this simulation, blood insulin was given as sine waves, and we examined the responses of each molecule. First, we simulated responses to a sine wave with an amplitude of 0.2 nM and a period of 120 minutes (Supp. Fig. 7A), mimicking additional secretion (Supp. Fig. 7B). In HFD mice, the responses of pAkt, pGSK3β, pFoxO1, and pACLY were drastically reduced by more than half. In contrast, the responses of pS6K and p4E-BP1 remained unchanged and were even larger than the reduction in IR expression. This indicates that pS6K and p4E-BP1 were unaffected by the reduction in IR or IRS2 expression levels, unlike other downstream molecules. Furthermore, we calculated the amplitudes of downstream molecules under various amplitudes and periods of blood insulin levels (Fig. 7A). Under both CD14w and HFD14w conditions, molecules with a small tau, such as pIR, pAkt, pGSK3β, and pFoxO1, responded to all secretion patterns (Fig. 3D). On the other hand, pACLY, which has a large tau, responded to additional and basal secretions, while pS6K and p4E-BP, which are regulated by an IFFL, were more responsive to additional secretion. These characteristics are consistent with our previous results[21]. In particular, p4E-BP1 responded to a narrower range of periods than pS6K, due to cooperative control (Hill equation). Furthermore, the responses of all molecules, including pIR, pAkt, pGSK3β, pFoxO1, and pACLY, were reduced under obese conditions (Fig. 7AB). However, despite the attenuation of the signal to pAkt under obese conditions, pS6K and p4E-BP1 showed almost the same response as the control (Fig. 7AB). This robustness might have been due to an IFFL, which was adjusted to enable pS6K and p4E-BP to respond to small changes in pAkt, thereby ensuring a stable response. In particular, the response of p4E-BP1 was even more stable under obese conditions due to the cooperative control by the Hill equation. These results indicate that the effects of obesity on molecular responses to physiological insulin secretion patterns varied between molecules. Specifically, pS6K and p4E-BP1, which are regulated by an IFFL, were less affected by obesity and reduced IR and IRSs expression, and thus maintained robust responses. These molecule-specific characteristics may contribute to selective insulin resistance.

**Fig. 7.**
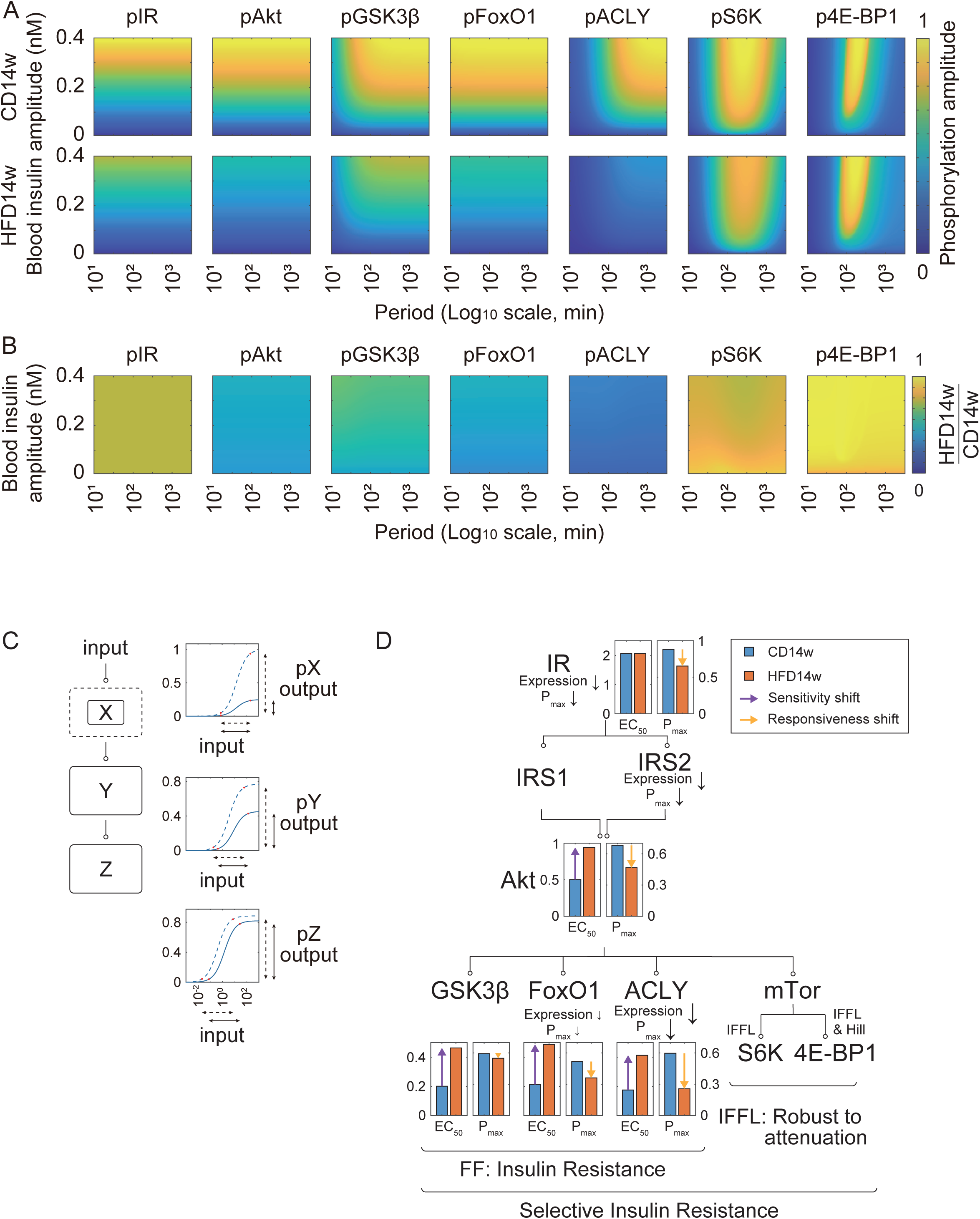
Selective insulin resistance is achieved through the characteristics of the insulin signaling pathway. (A) The response of each molecule at varied amplitudes and periods. The peak values of CD14w and HFD14w mice are shown in the color plots. The x-axis indicates amplitude on a logarithmic scale. (B) HFD14w mice to CD14w mice ratios of the mean time variability of each molecule are shown in the color plots. The x-axis indicates amplitude on a logarithmic scale. (C) Characteristics of the signaling pathway revealed in this study. Changes in signal transduction when the expression level of X is reduced (from dotted to solid line) in a hypothetical signaling pathway consisting of Input, X, Y, and Z are shown. The dynamic range of the input and output is defined as the range over which the signal varies from 5% to 95% of its maximum. When the expression of X decreases, the input dynamic range remains unchanged (x-axis), whereas the output dynamic range narrows (y-axis). In molecule Y, the input dynamic range shifts to the right, and the output dynamic range is partially restored. Likewise, in molecule Z, the input dynamic range shifts further to the right, and the output dynamic range recovers further. (D) Summary of the mechanisms underlying the alteration in the insulin signaling pathway molecules during obesity progression revealed in this study. The EC_50_ and P_max_ values for each molecule are shown as dotted (CD14w) and solid (HFD14w) bar graphs. The EC_50_ and P_max_ values are indicated in dark blue and yellow, respectively. All molecules with feedforward (FF) regulation exhibited an increase in their EC_50_. Although the P_max_ is restored along with the signaling cascade, the P_max_ of ACLY decreased because its protein expression levels decreased. In contrast, pS6K and p4E-BP1 maintained robust responses despite the reduction in the P_max_ of Akt due to their network structure, incoherent-feedforward-loop (IFFL) regulation.

## 4 Discussion

Insulin resistance, which attenuates the hypoglycemic effect of insulin, has been proposed to be characterized by two aspects, the sensitivity shift and the responsiveness shift, corresponding to a rightward shift in the dose–response curve and a decrease in the maximum value, respectively[2,19,20]. In this study, the sensitivity and responsiveness shifts correspond to an increase in the EC_50_ and a decrease in the P_max_, respectively. Our experiments and mathematical analysis revealed that decreases in the expression levels of upstream molecules (decreases in their P_max_) propagated to affect both the EC_50_ and P_max_ of downstream molecules via characteristics of FF (Figs. 4A–C, 7C, and Supp. Fig. 5A). However, due to the trade-off relationship between the EC_50_ and P_max_ (Supp. Fig. 5B), the decreased P_max_ of an upstream molecule recovered as the EC_50_ of downstream molecules increased. Thus, the insulin signaling pathway may prioritize maintaining the responsiveness of downstream molecules over regulating the EC_50_. This may be because, while the biological system can compensate for an increased EC_50_ by hyperinsulinemia, restoration of the P_max_ requires increasing the expression levels of signaling molecules, which may entail a high biological cost and complex mechanisms. In actuality, insulin demand and fasting blood insulin levels showed a high correlation, indicating compensation for the response (Fig. 5CD). These results suggest that the increase in the EC_50_ and the compensatory increase in insulin secretion may form a vicious cycle that exacerbates insulin resistance.

Numerous previous studies have investigated IR in the context of insulin resistance. Mutations in IR associated with insulin resistance have been identified in humans[3,4]. There are reports that even at a 2.4% reduction in IR expression, glucose oxidation remains comparable to normal levels[12] and that IR heterozygous knockout mice show no differences in Akt phosphorylation and glucose metabolism[13]. These previous reports suggested that a decrease in IR expression may not necessarily affect downstream molecules (responses), thereby complicating our understanding of the contribution of IR expression changes to insulin resistance. The characteristics identified in this study can explain these previous experimental results (Supp. Fig. 6C) and indicate that the characteristics of the signaling pathway itself underlie these complicated phenotypes. Thus, understanding insulin resistance at the level of signal transduction requires consideration of the underlying information-processing mechanisms.

Blood insulin exhibits several temporal patterns, which have been shown to be important for insulin action[33–36]. Furthermore, our previous studies on rats revealed that downstream molecules in the insulin signaling pathway can be selectively regulated depending on the temporal patterns of insulin[21]. In this study, we confirmed that similar characteristics are conserved in mice, indicating that this characteristic transcends species. Interestingly, in HFD mice exhibiting insulin resistance, qualitative molecular responses, including sustained responses of pIR, pAkt, pGSK3β, pFoxO1, and pACLY and transient responses of pS6K and p4E-BP, were maintained. However, the quantitative responses varied depending on the molecule (Figs. 3, 4C, and 7B). Notably, ACLY phosphorylation levels in HFD mice were reduced to approximately one-third of those in control mice. This reduction was primarily attributed to reduced ACLY expression (Figs. 5A and 7D). ACLY expression decreased by the third week after obesity induction, consistent with previous reports showing a 75% reduction in liver ACLY by the fourth week after obesity induction[26]. ACLY catalyzes the conversion of citrate supplied from mitochondria to acetyl-CoA[37,38]. In this study, insulin resistance was induced by feeding mice an HFD. Therefore, a responsiveness shift might have occurred to address the increase in free fatty acids due to enhanced fat uptake before the attenuation of insulin signaling. GSK3β, a kinase that inhibits glycogen synthase, is inactivated by its phosphorylation via Akt[39,40]. In our experiment, GSK3β did not show an expression change but exhibited only a sensitivity shift. This may be because glycogen synthesis needs to be maintained even under the obese condition. FoxO1 is a transcription factor related to gluconeogenesis[40,41]. FoxO1 exhibited decreased expression and both sensitivity and responsiveness shifts. These changes are considered to be one of the causes of insulin resistance observed in the IPITT experiments, as they attenuate the reduction in blood glucose levels (Supp. Fig. 1C–F). The responsiveness shift caused by a decrease in IR expression is restored by the characteristics of the signaling pathway; however, this is offset by the sensitivity shift. As described above, the responsiveness shift is further modulated by the expression levels of downstream molecules. Thus, insulin sensitivity is thought to be individually adjusted for each pathway.

On the other hand, the responses of pS6K and p4E-BP1 in HFD mice remained comparable to those in CD mice (Fig. 7AB). This is attributed to the IFFL, which responds to changes in concentration. Although insulin secretion induced by meals is unlikely to exhibit changes as abrupt as the pulse stimulation used in this study, a robust response is thought to be ensured by parameters that allow the IFFL to respond even to small changes; in fact, pS6K and p4E-BP1 can respond to a 120-minute sinusoidal wave, which mimics additional secretion. Although a negative feedback loop can account for these characteristics, we selected the IFFL based on our previous studies[21,25]. While the exact network structure remains unclear, the overall conclusion is expected to remain unchanged. S6K is a protein synthesis-related factor[40,42], and robust responses are likely required for postprandial protein synthesis even under obese conditions. In the Mouse Insulin Signaling Model, p4E-BP1 exhibited a more robust response to obesity than pS6K due to its cooperative response, as described by the Hill equation. These results suggest that 4E-BP1 plays an important role in insulin action in the liver during obesity progression. The phosphorylation-mediated regulation of 4E-BP1 has been reported to involve initial phosphorylation at T37/T46, resulting in partial structural changes, followed by phosphorylation at S65/T70 for further stabilization[24]. These multi-step phosphorylation and structural changes may underlie the switch-like response of 4E-BP1 to pAkt observed in this study (Fig. 2A). Our Mouse Insulin Signaling Model incorporates this mechanism using the Hill equation; however, further studies are needed to explain the mechanism in detail. When 4E-BP1 is phosphorylated by mTOR, it dissociates from eIF4E, a cap-binding protein, and promotes translation[43]. 4E-BP1/2 double-knockout mice exhibit increased lipid accumulation in the liver under HFD conditions compared to wild type mice[44]. Additionally, eIF4E +/- mice maintain normal protein synthesis but show suppressed lipid accumulation in the liver and do not develop obesity even under HFD conditions[45]. Thus, 4E-BP1 may regulate obesity-related responses and may be essential for maintaining robust responses under obese conditions. Taken together, these results suggest that 4E-BP1 may contribute to selective insulin resistance, where gluconeogenesis is impaired but lipid accumulation is enhanced in the liver. As a result, many aspects of insulin resistance, including metabolic responses, could be explained by alterations in the insulin signaling pathway.

The spare receptor hypothesis has been proposed for many receptors[20,46], including the IR; even a small amount of ligand binding can achieve a maximum response. The significance of spare receptors is that they may confer robustness against receptor loss due to disease and ensure linearity with respect to ligands[15]. In our analysis, we found that as IR expression decreased, the changes in the sensitivity shift (EC_50_), responsiveness shift (P_max_), and time constant (tau) of downstream molecules increased (Fig. 4D and Supp. Fig. 5D), thereby ensuring a robust response of the insulin signaling pathway when IR is excessively expressed. Furthermore, we examined the significance of excessive IR receptors by reducing EC_50_ in simulations (Fig. 6BC). When the EC_50_ of IR was lowered and other parameters were adjusted to maintain the EC_50_ of Akt, the P_max_ of Akt decreased. While increased expression of Akt can restore its P_max_, it would incur additional biological costs. In other words, while lowering the EC_50_ of IR may reduce the required IR expression level, increasing the expression levels of downstream molecules is necessary to ensure the overall response of the signaling pathway. Given that signaling pathways branch out further downstream, excessive expression of upstream receptors may be the most effective strategy to minimize the biological cost of the signaling pathway. Thus, one of the reasons for the existence of spare receptors is that they enable efficient and robust signal transduction. Additionally, P_max_ represents the dynamic range of the output, and its reduction leads to a shift in the responsiveness of downstream molecules; however, this cannot be achieved solely by increasing the number of spare receptors, but rather requires coordinated regulation throughout the entire signaling pathway. The propagation of the effects of differences in upstream molecule expression to downstream molecules may help explain individual differences in insulin sensitivity. Additionally, treatments for insulin resistance caused by reduced IR expression may involve restoring the expression levels of the depleted molecules; this could serve as a comprehensive therapeutic approach that restores not only the response of the insulin signaling pathway but also hyperinsulinemia. The characteristics and mechanisms revealed in this study, including those related to the “spare receptor hypothesis,” are considered to be general features conserved across signaling pathways. Many diseases involve altered receptor expression, and such changes are thought to induce the sensitivity and/or responsiveness shifts in downstream molecules. A detailed grasp of these mechanisms may provide a deeper understanding of disease pathogenesis and potential treatments.

Negative feedback regulation via IRS has been reported for the insulin signaling pathway[42]. On the other hand, experiments using transgenic mice and inhibitors have challenged the existence of such negative feedback regulation[47,48]. According to these studies, the presence of this feedback in vivo is controversial. The Mouse Insulin Signaling Model simplifies detailed processes such as IR internalization. Furthermore, the insulin signaling pathway is influenced by various factors, including lipids in the blood and hormones such as adipokines. The insulin signaling pathway is highly conserved across organs and species. In this study, we performed experiments using murine liver; however, it remains unclear whether obesity-induced alterations are conserved in other organs or across species. We observed that similar dynamic properties of molecules in the insulin signaling pathway are also maintained in rat liver[21], suggesting interspecies conservation. Furthermore, the characteristics we identified, including those of the “spare receptor hypothesis”, represent fundamental properties of signaling pathways. In other words, even if the regulation of the insulin signaling pathway described above may differ to some extent among organs or species, the underlying properties we discovered appear to be conserved. Further studies using models developed based on our model would be necessary to deepen our understanding of the contributions of other molecules and clarify how they affect the insulin signaling pathway. In particular, assessing the dynamic properties of molecules in human organs remains a big challenge. Nevertheless, based on the model we developed in this study, we believe that understanding the dynamic properties of insulin responses in humans, as well as their alterations in obesity, will also be advanced.

Our Mouse Insulin Signaling Model describes reactions using mass action kinetics without considering enzyme–substrate complexes. However, many studies, including our previous models, have also employed mass action kinetics without considering enzyme–substrate complexes to describe the reactions of signal transduction pathways and have generated numerous findings using this approach[21,25,49]. Therefore, the characteristics and mechanisms identified in this study are considered to be conserved across many signaling pathways. On the other hand, some biological reactions, such as zero-order ultra-sensitivity[50], oscillation dynamics[51,52] and n-th order reaction[53], require the consideration of enzyme–substrate intermediates and/or coordinate responses, and thus may need to be examined individually under specific contexts.

Despite these limitations, our study revealed novel properties of the signaling pathway and provided new insights into the mechanisms underlying the “spare receptor hypothesis.” Furthermore, it provides a unified and quantitative framework for understanding insulin resistance. Our findings are expected not only to advance the mechanistic understanding of insulin resistance and its therapeutic applications but also to broaden our understanding of other signaling pathways that exhibit spare receptor properties.

## CRediT authorship contribution statement

**So Morishita:** Writing – review & editing, Writing – original draft, Visualization, Validation, Software, Methodology, Investigation, Funding acquisition, Formal analysis, Data curation, Conceptualization. **Shinsuke Uda:** Writing – review & editing, Validation, Supervision, Formal analysis. **Hiroyuki Kubota:** Writing – review & editing, Writing – original draft, Validation, Supervision, Resources, Project administration, Methodology, Funding acquisition, Formal analysis, Conceptualization.

## Acknowledgments

The infrastructure of the Omics Science Center Secure Information Analysis System, Medical Institute of Bioregulation at Kyushu University provided computational resources.

## Data and material availability

Any additional information required to reanalyze the data reported in this paper is available from the lead contact upon reasonable request.

## Supplementary figure legends

**Supp. Fig. 1.**
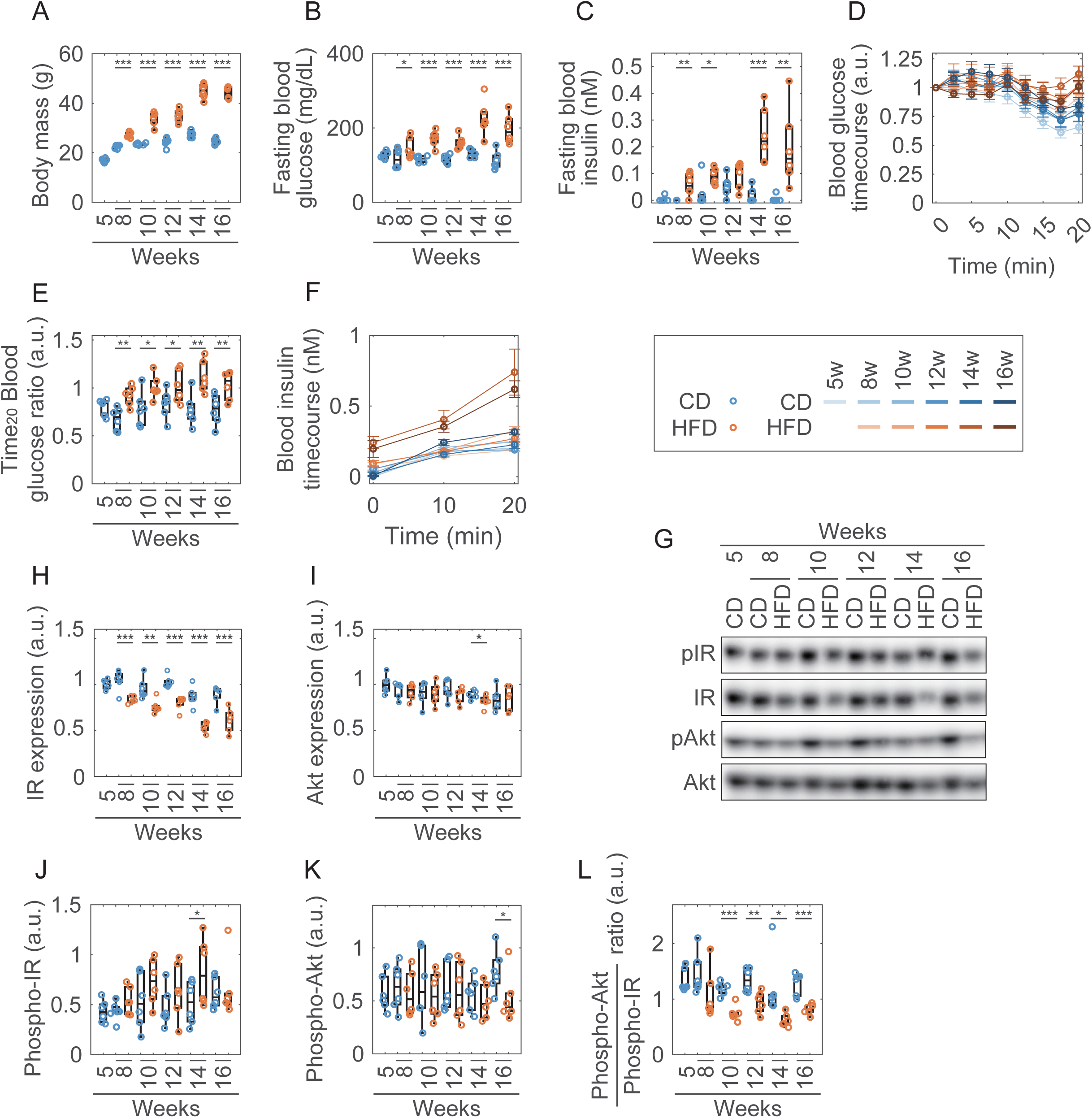
Week-of-age-dependent changes in physiological data and molecular expressions, related to Fig. 1. (A) Body mass; (B) blood glucose; and (C) blood insulin levels of overnight-fasted mice under the indicated conditions. (D) Time courses of blood glucose levels during the intraperitoneal insulin tolerance test (IPITT). The values are normalized to those at 0 minutes. (E) Ratio of blood glucose levels at 20 minutes to those at 0 minutes in the IPITT. (F) Time courses of blood insulin levels during the IPITT. (G) Representative Western blot images of pIR, IR, IRS1, IRS2, pAkt, and Akt in the liver at 20 minutes. (H) Protein expression level of IR; (I) protein expression level of Akt; (J) phosphorylation level of IR; (K) phosphorylation level of Akt; and (L) their ratio (pAkt/pIR). Data are shown as mean ± SD (n=6). *** p < 0.001, ** p < 0. 01, * p < 0.05, Welch’s *t*-test.

**Supp. Fig. 2.**
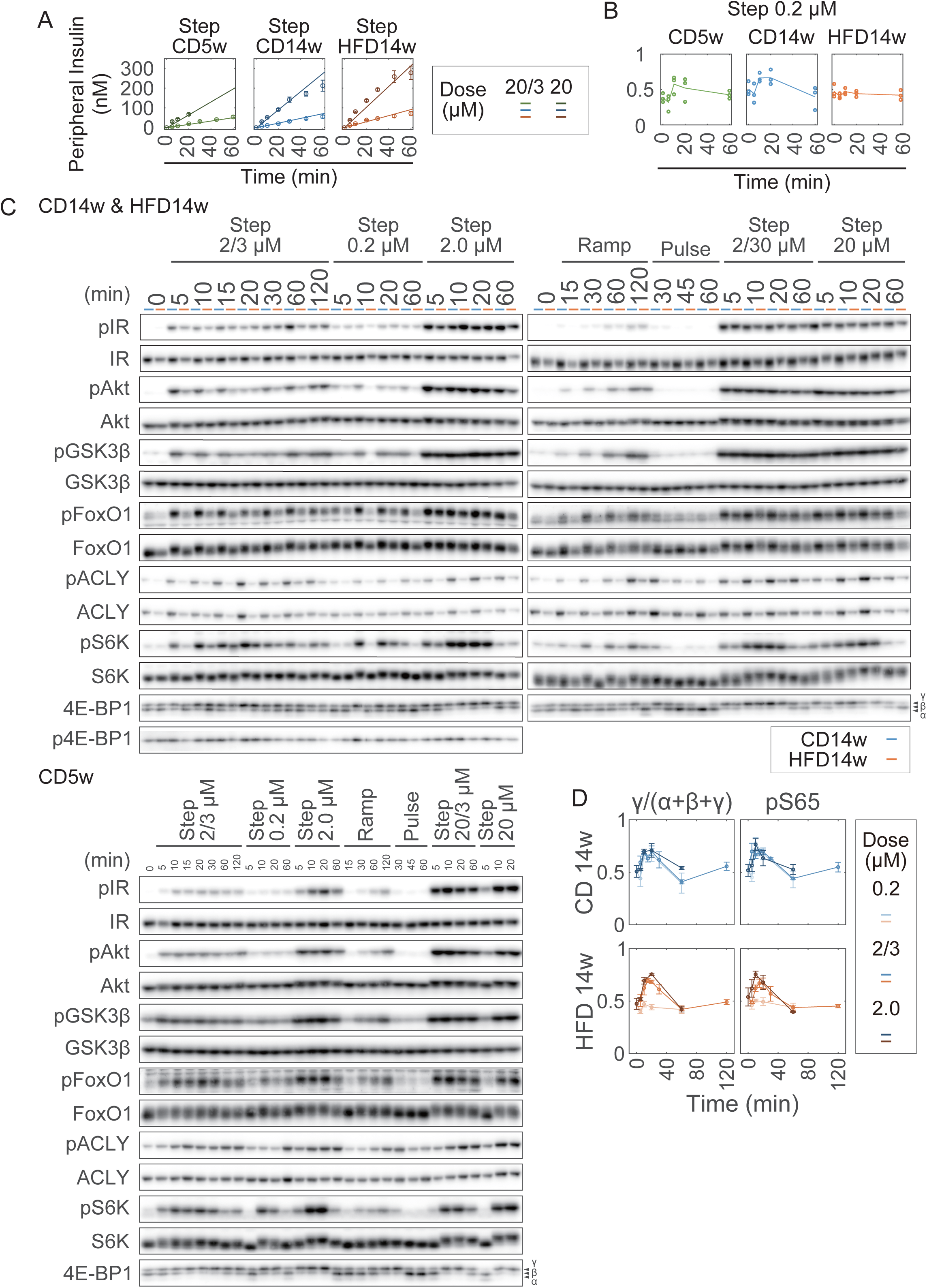
Temporal profiles of blood insulin, pIR, pAkt, pGSK3β, pFoxO1, pACLY, pS6K, and 4E-BP1 in the liver, related to Fig. 2. (A) Time courses of blood insulin levels under high-dose insulin stimulation. Solid lines and dots indicate simulation results and experimental data, respectively. (B) p4E-BP1 during the infusion experiment of step 0.2 µM stimulation. Lines and dots indicate the mean and experimental data, respectively. (C) Representative Western blot images corresponding to Fig. 1F. (D) Comparison of band shift ratio (γ/(α+β+γ)) and pSer65 of 4E-BP1. Data are shown as mean ± SD (n=3).

**Supp. Fig. 3.**
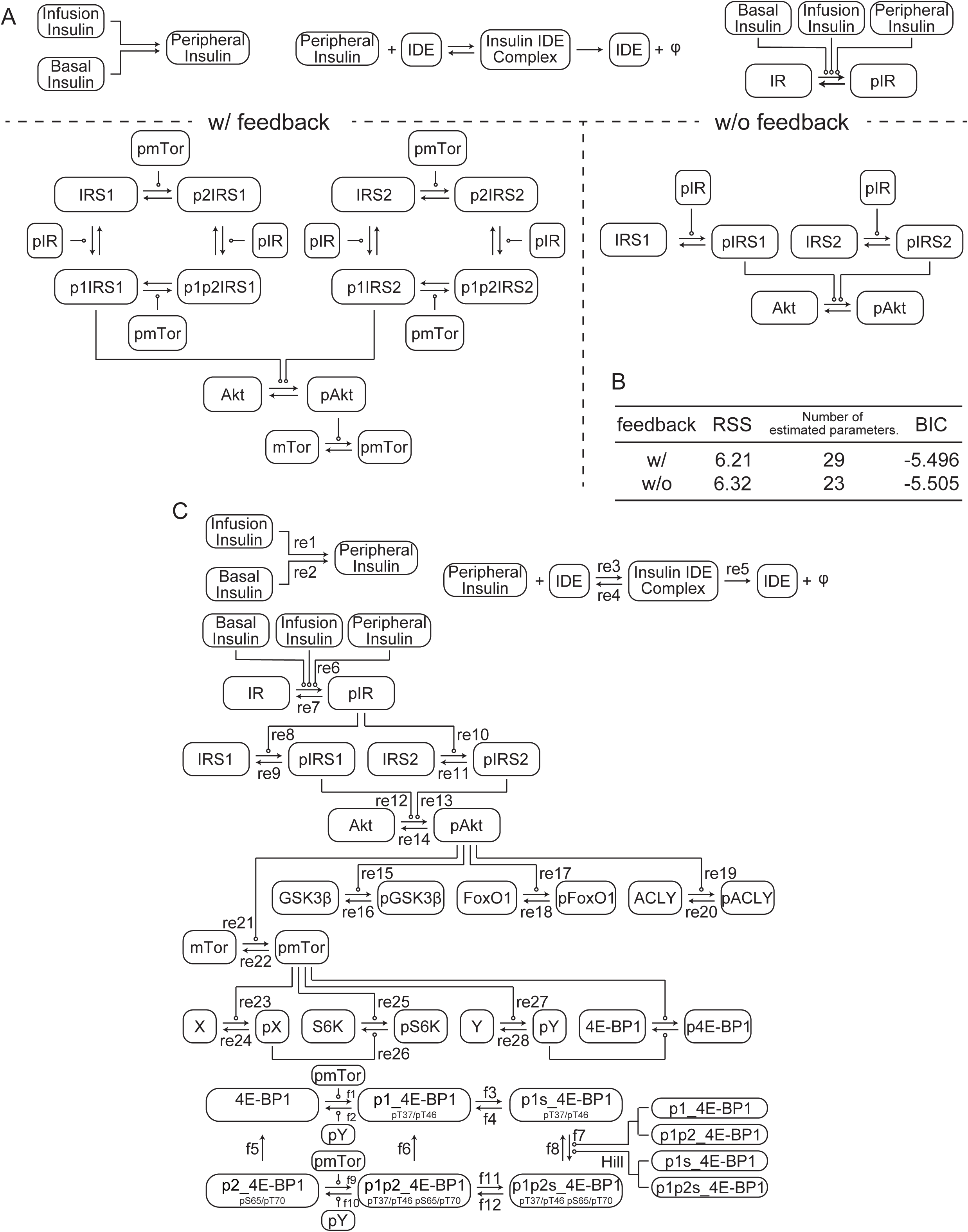
The insulin signaling pathway model, related to Fig. 3. (A) Schematic overview of feedback from mTOR to IRSs for comparison. (B) Comparison of models with and without feedback from mTOR to IRSs (see Star Methods). RSS and BIC represent the residual sum of squares and Bayesian information criterion, respectively. For BIC, a smaller value indicates a better model. (C) Schematic overview of the insulin signaling pathway. “re#” and “f#” indicate the reaction number (Supp. Tables 1 and 3) and the flux number (Supp. Tables 1–3), respectively. IDE represents insulin-degrading enzyme.

**Supp. Fig. 4.**
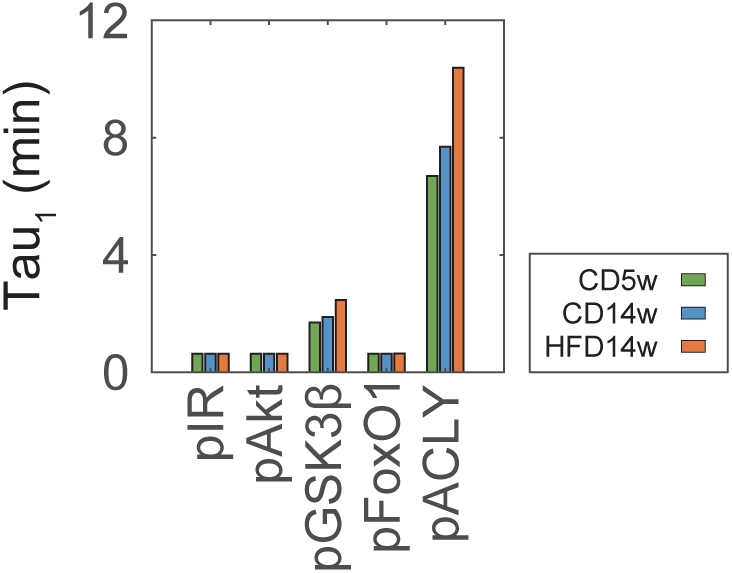
Tau at a blood insulin concentration of 1 nM (Tau_1_), related to Fig. 3. Tau at 1 nM blood insulin concentration (Tau_1_).

**Supp. Fig. 5.**
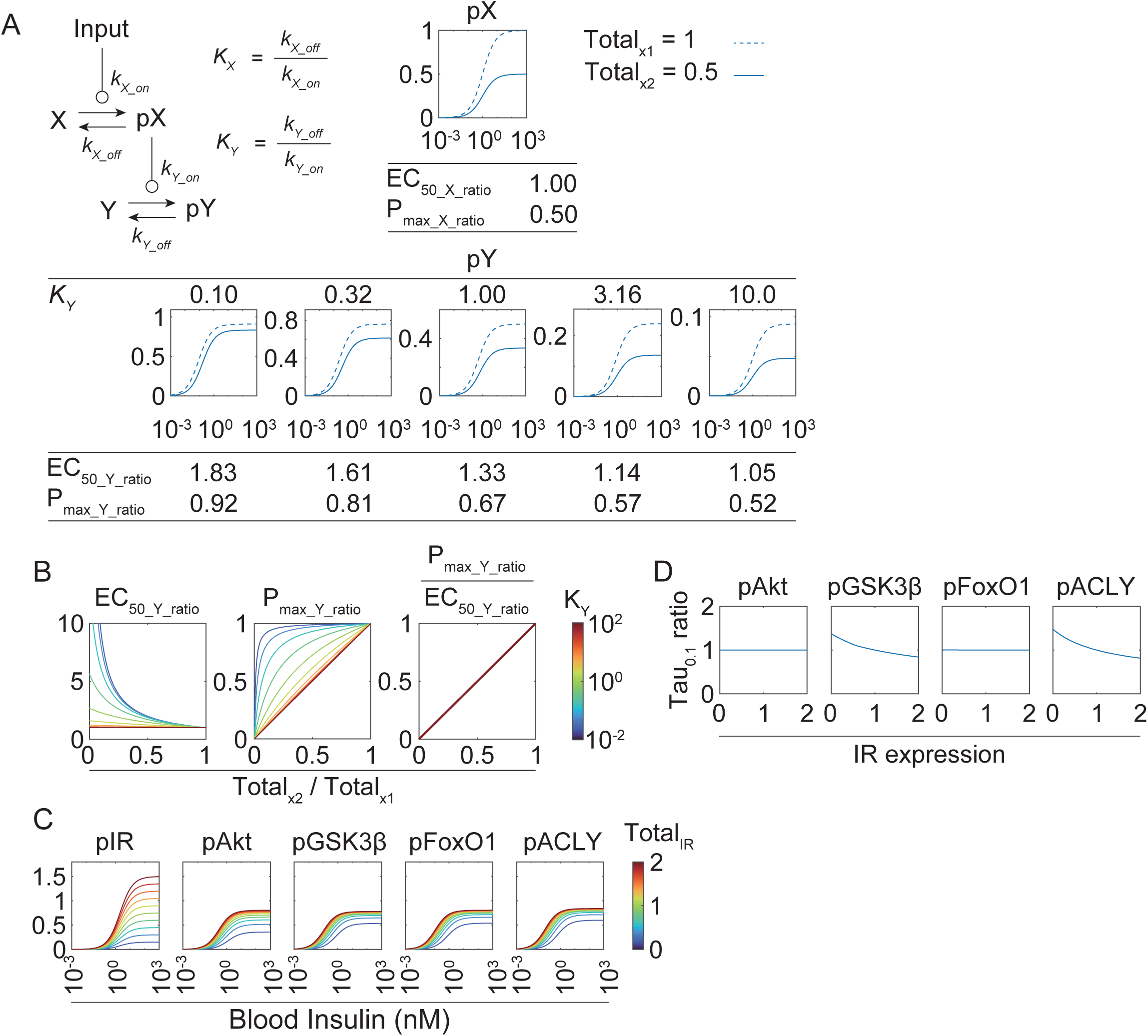
Characteristics of receptor-expression-dependent signaling pathway, related to Fig. 4. (A) Characteristics of the signaling pathway revealed by the toy model. Changes in the EC_50_ and P_max_ were analyzed when the expression level of X (*Total_X_*) was halved in a hypothetical signaling pathway consisting of Input, X, and Y. The K_y_-dependent dose–response curve of pY and the corresponding values of EC_50_Y_ratio_ (EC_50_Y2_/EC_50_Y1_) and P_max_Y_ratio_ (P_max_Y2_/ P_max Y1_) are shown. Solid and dotted lines represent the results for *Total_X_* = 1 and *Total_X_* = 0.5, respectively. See Star Methods for details. (B) Protein expression ratio of X (*Total_X2_*/*Total_X1_*)-dependent EC_50_Y_ratio_ (left) and of P_max_Y_ratio_ of Y (middle), and their ratio (P_max_Y_ratio_/ EC_50_Y_ratio_ [right]). (C) Change in the dose–response curves of downstream molecules with varying IR expression level. The rate constants obtained through parameter estimation were used to calculate the changes in the dose responses of downstream molecules as IR expression levels were varied from 0.1- to 2-fold. (D) Changes in the Tau_0.1_ ratio of downstream molecules in response to the changes in IR and Akt expression levels. The rate constants obtained through parameter estimation were used to calculate the changes in Tau_0.1_ of downstream molecules. The Tau_0.1_ values of CD14w were normalized to 1.

**Supp. Fig. 6.**
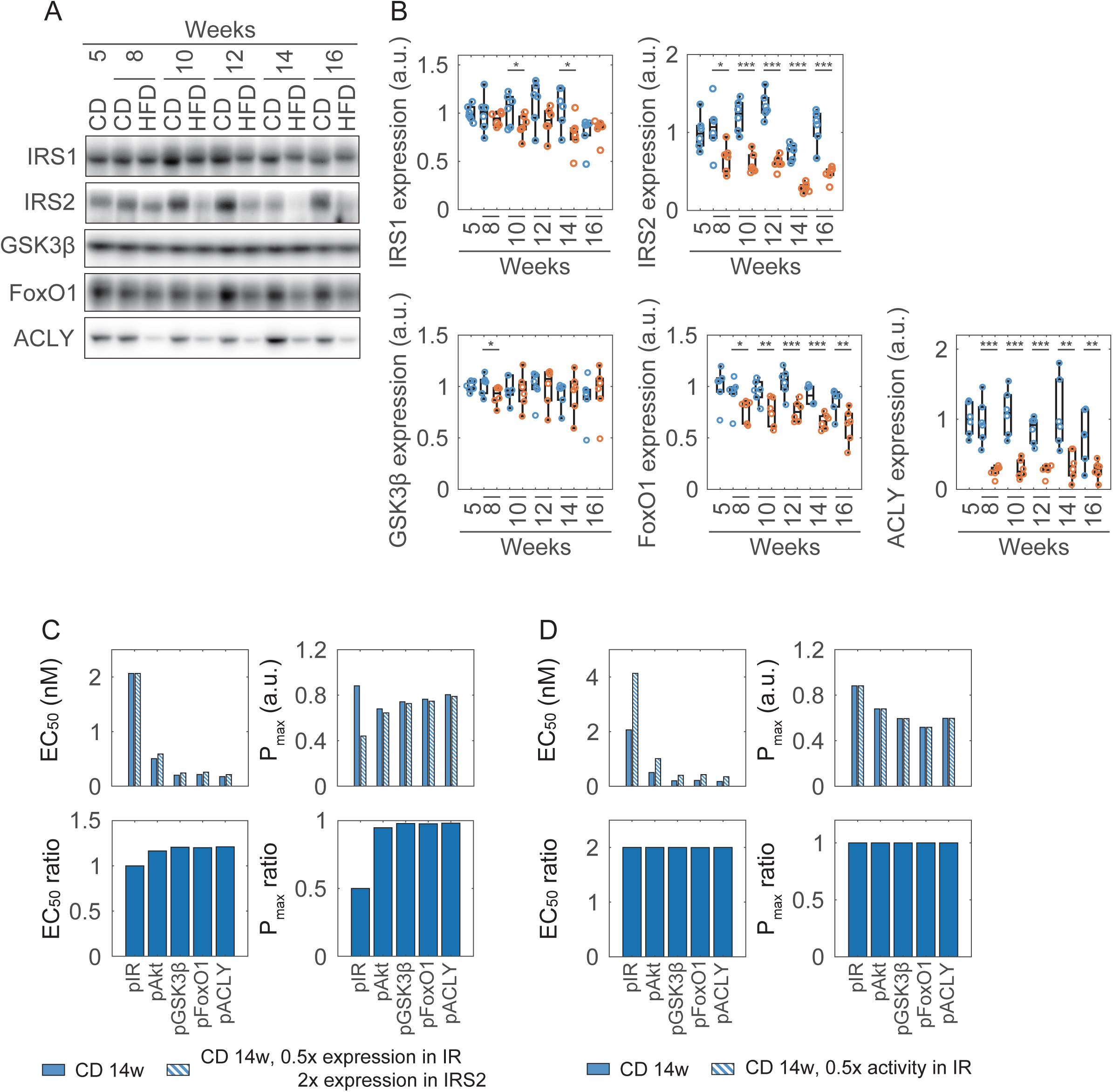
Temporal profiles of GSK3β, FoxO1, and ACLY expression levels and simulation results under conditions of the previous study, related to Fig. 5. (A) Representative Western blots of the indicated molecules; and (B) their expression levels in the liver at 20 minutes. Data are shown as mean ± SD (n = 6). *** p < 0.001, ** p < 0.01, * p < 0.05, Welch’s *t*-test. (B) Reproduction of the results of Merry et al. (2017). The EC_50_ and P_max_ of downstream molecules were calculated under the original CD14w condition and under the condition described by Merry et al., in which IR expression is halved and IRS2 expression is doubled. The ratios between conditions are shown at the bottom as the EC_50_ ratio and P_max_ ratio. (C) The EC_50_ and P_max_ of downstream molecules were calculated under the original CD14w condition and under the condition of halved IR phosphorylation activity. The ratios between them are shown at the bottom as EC_50_ ratio and P_max_ ratio.

**Supp. Fig. 7.**
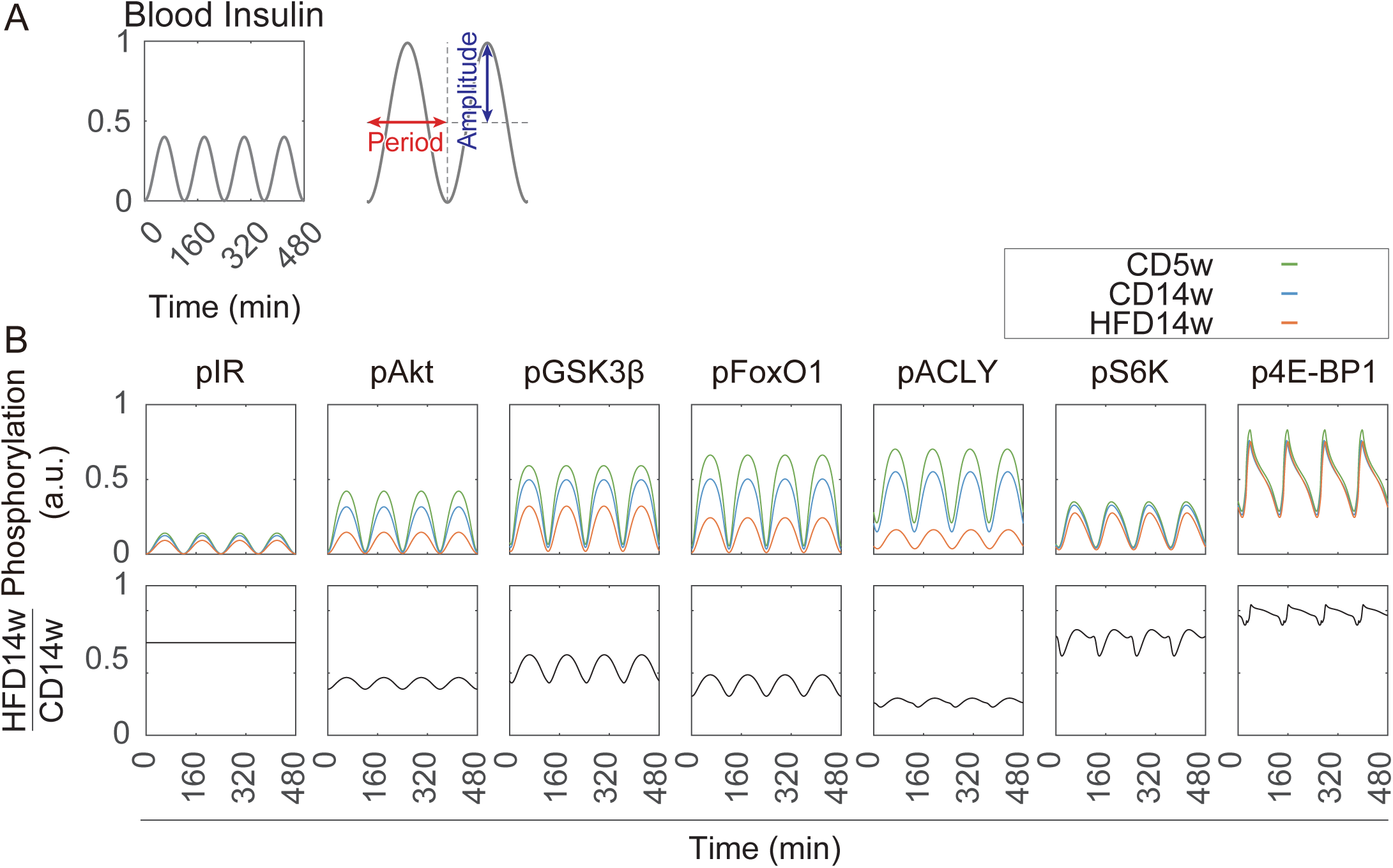
Responses to a sinusoidal wave of blood insulin, related to Fig. 7. (A) Example of a sine wave of blood insulin levels, resembling an additional secretion. Amplitude: 0.2 nM; period: 120 minutes. (B) Responses of each molecule to the sine wave of blood insulin levels with an amplitude of 0.2 nM and a period of 120 minutes. The upper panel shows the values for CD5w (green), CD14w (blue), and HFD14w (red). The lower panel shows the ratio of CD14w to HFD14w.

## Supplementary table legends

**Supp. Table 1.**
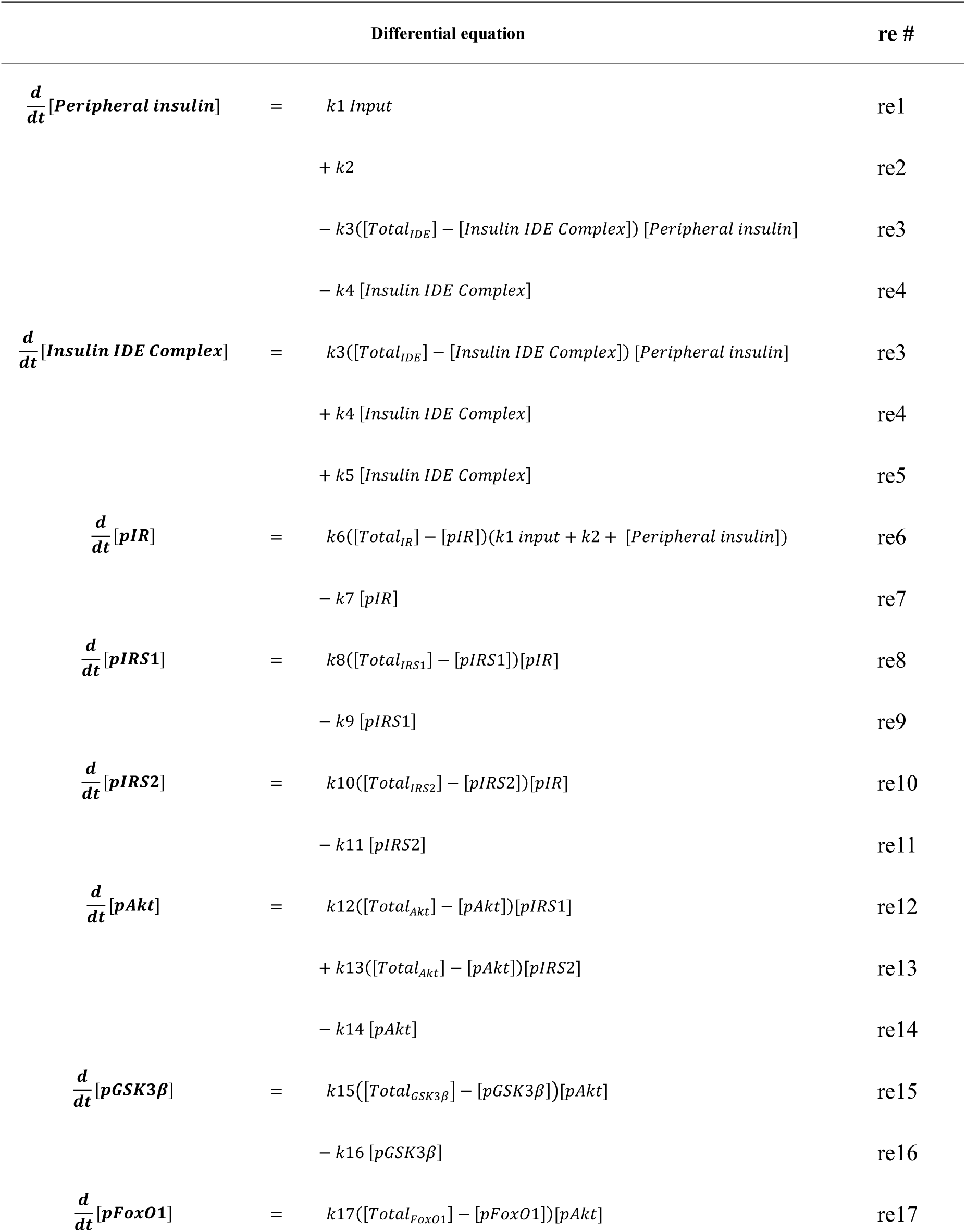
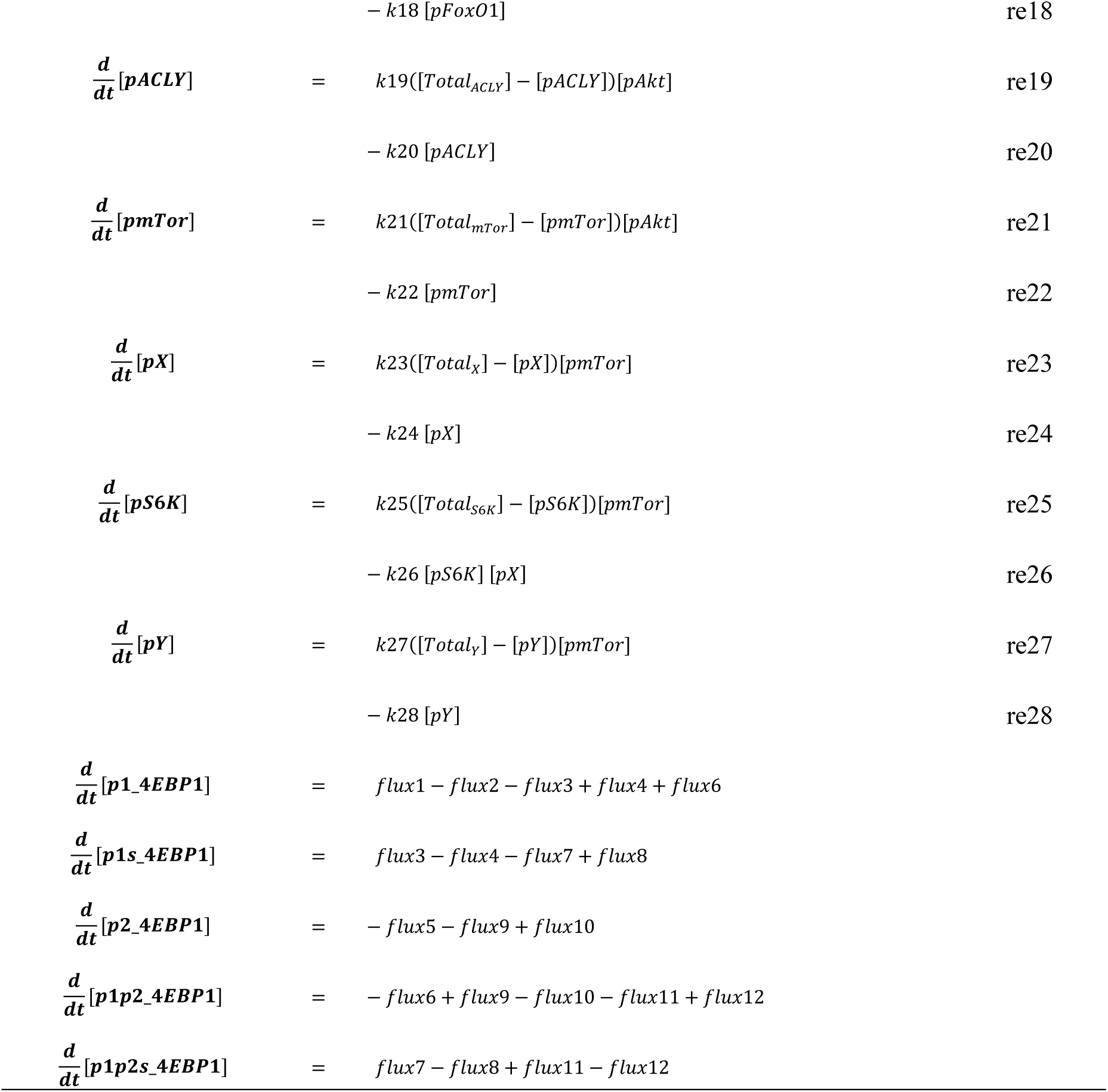
Differential equations of the insulin signaling pathway model, related to Fig. 3. Ordinary differential equations (ODEs) of the insulin signaling pathway model. “re#” and “f#” indicate reaction number and flux number (Supp. Fig. 3, Supp. Tables 2 and 3), respectively. 4E-BP1 fluxes are shown in Supp. Fig. 6. The simulation was performed using MATLAB (see Method Details under Star Methods).

**Supp. Table 2.**
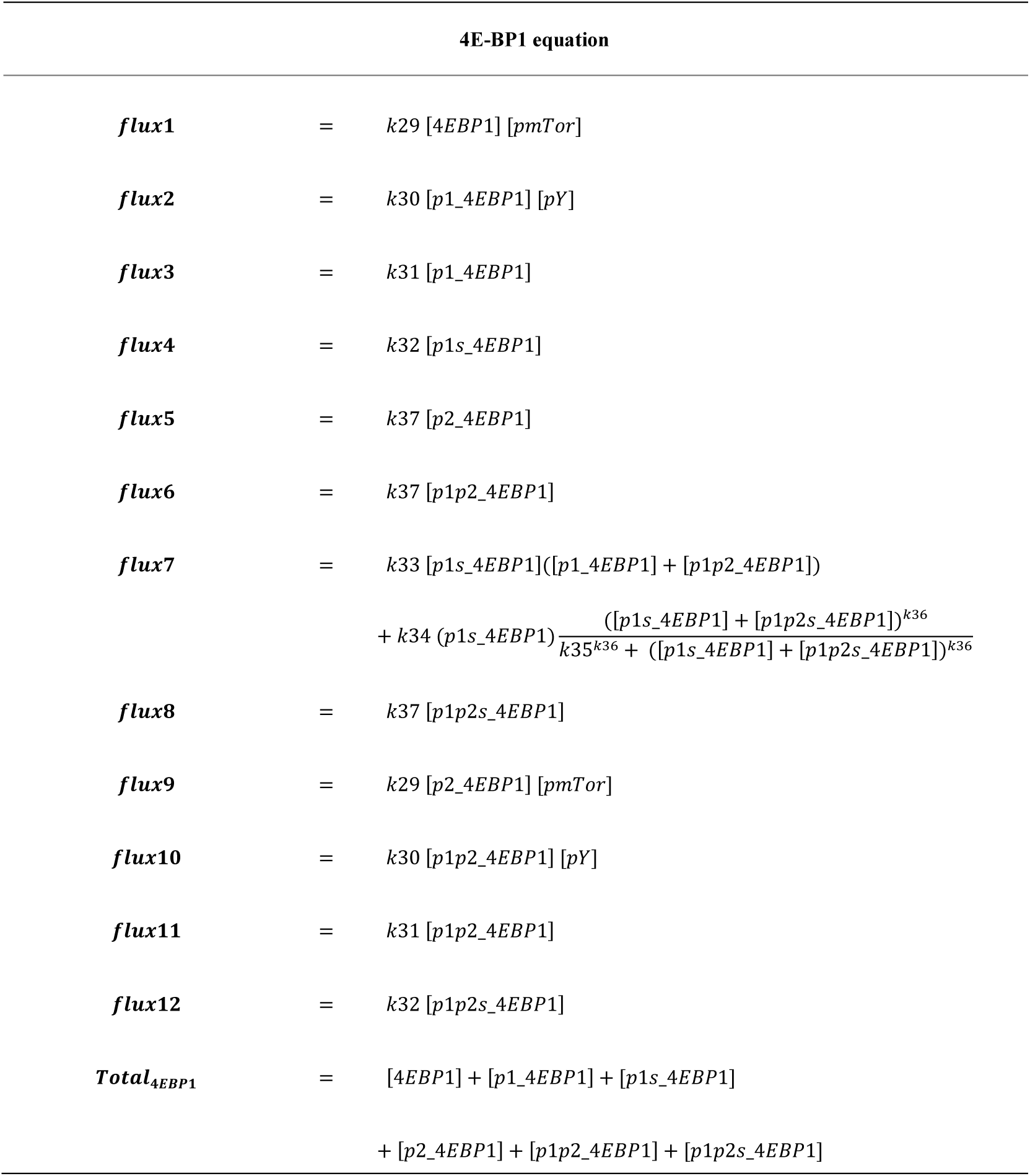
Differential equations of 4E-BP1, related to Fig. 3. 4E-BP1 fluxes in ordinary differential equations (ODEs) (Supp. Table 1).

| 4E-BP1 equation |  |  |
| --- | --- | --- |
| *flux1* | = | $k_{29} [4EBP1] [pmTor]$ |
| *flux2* | = | $k_{30} [p1\_4EBP1] [pY]$ |
| *flux3* | = | $k_{31} [p1\_4EBP1]$ |
| *flux4* | = | $k_{32} [p1s\_4EBP1]$ |
| *flux5* | = | $k_{37} [p2\_4EBP1]$ |
| *flux6* | = | $k_{37} [p1p2\_4EBP1]$ |
| *flux7* | = | $k_{33} [p1s\_4EBP1]([p1\_4EBP1] + [p1p2\_4EBP1])$<br>$+ k_{34} (p1s\_4EBP1) \frac{([p1s\_4EBP1] + [p1p2s\_4EBP1])^{k_{36}}}{k_{35}^{k_{36}} + ([p1s\_4EBP1] + [p1p2s\_4EBP1])^{k_{36}}}$ |
| *flux8* | = | $k_{37} [p1p2s\_4EBP1]$ |
| *flux9* | = | $k_{29} [p2\_4EBP1] [pmTor]$ |
| *flux10* | = | $k_{30} [p1p2\_4EBP1] [pY]$ |
| *flux11* | = | $k_{31} [p1p2\_4EBP1]$ |
| *flux12* | = | $k_{32} [p1p2s\_4EBP1]$ |
| *Total<sub>4EBP1</sub>* | = | $[4EBP1] + [p1\_4EBP1] + [p1s\_4EBP1]$<br>$+ [p2\_4EBP1] + [p1p2\_4EBP1] + [p1p2s\_4EBP1]$ |

**Supp. Table 3.**
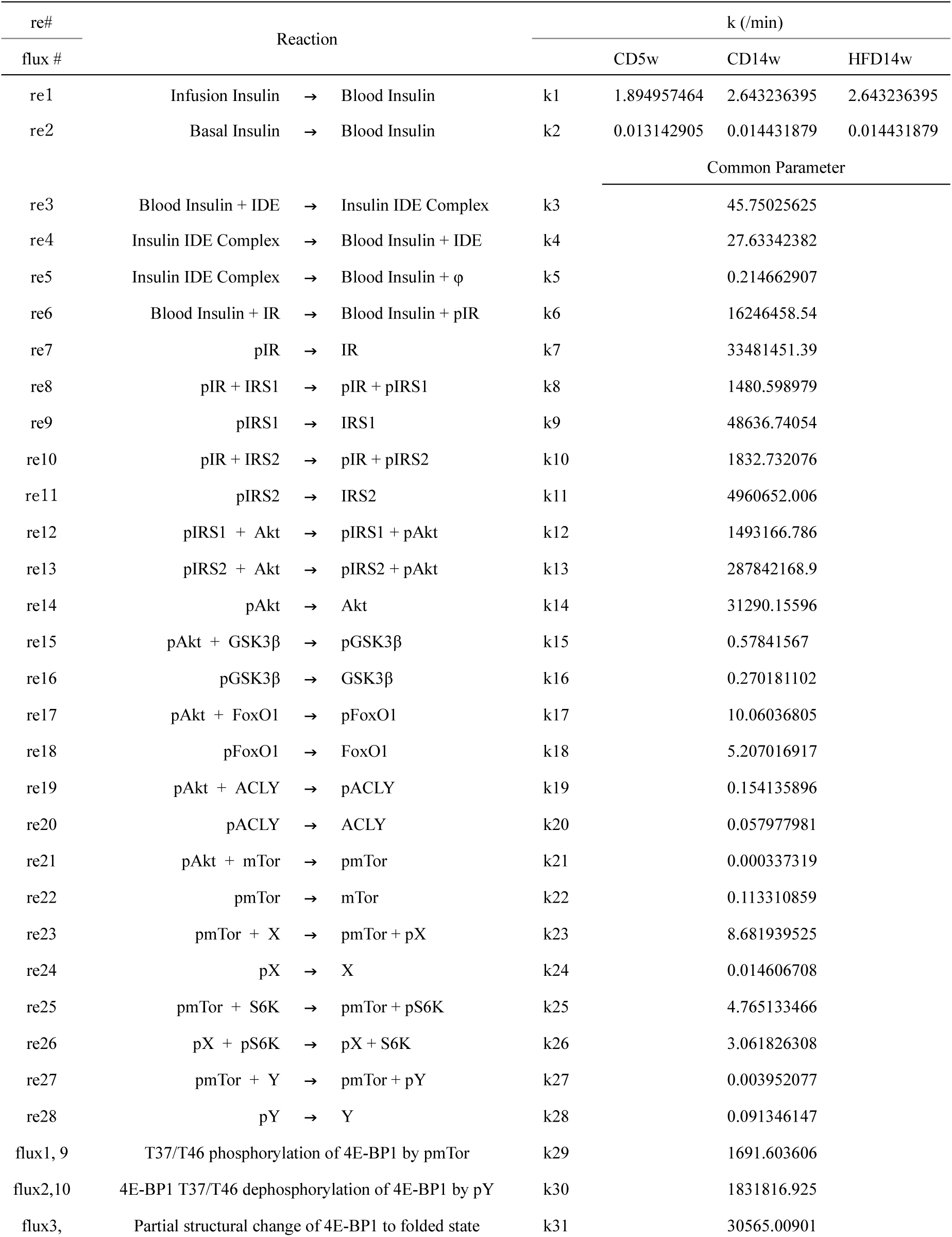
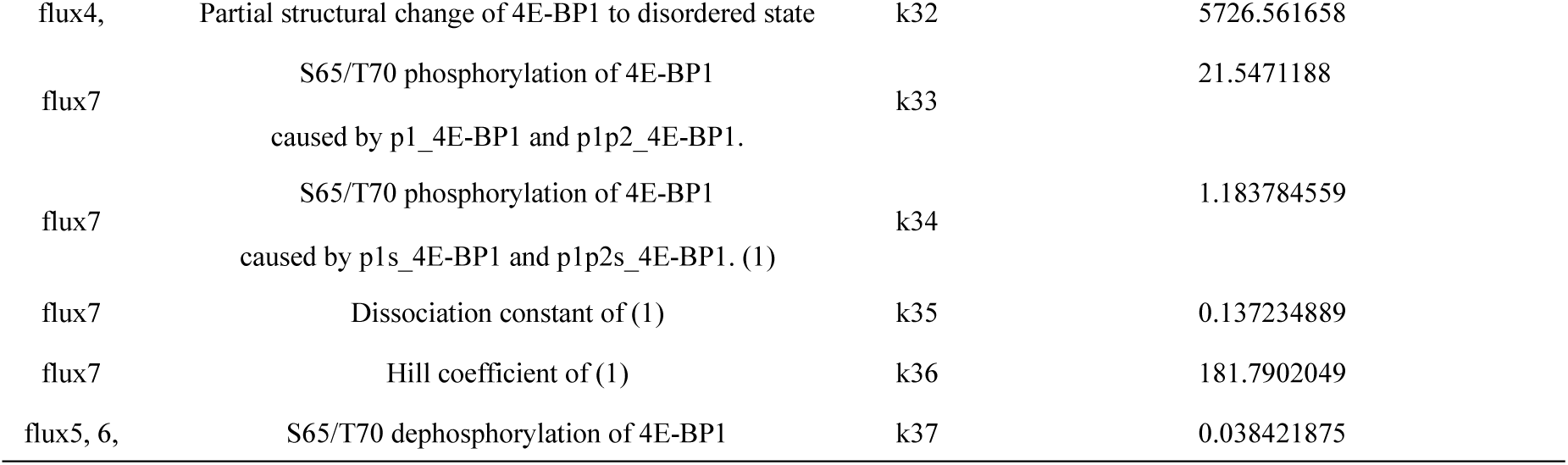
Reactions and parameters of the insulin signaling pathway model, related to Fig. 3. Reactions and rate constants in the insulin signaling pathway model. “re#” and “f#” indicate the reaction number and the flux number (Supp. Fig. 3, Supp. Tables 1 and 2). The simulation was performed using MATLAB (see Method Details under Star Methods).

**Supp. Table 4.** Total and initial parameters of the insulin signaling pathway model, related to Fig. 3. Total amount, initial amount (peripheral insulin, insulin degradation enzyme complex), and initial phosphorylated amount (signaling pathway molecules) in the insulin signaling pathway model. Values indicated in red are relative values (fold changes) compared to CD5w protein expression levels.

| molecule | Common parameter model |  |  |  |  |  |
| --- | --- | --- | --- | --- | --- | --- |
|  | Total amount |  |  | Initial (phosphorylated) amount |  |  |
|  | CD5w | CD14w | HFD14w | CD5w | CD14w | HFD14w |
| Peripheral insulin |  |  |  | 0.021144 | 0.017129 | 0.049827 |
| IDE Complex | 1.823768 | 2.456373 | 1.59704 | 0.061226 | 0.06723 | 0.120839 |
| IR | 0.850376 | 0.749933 | 0.55685 | 0.013917 | 0.011312 | 0.019746 |
| IRS1 | 1 (fixed) | 1.059855 | 0.783604 | 0.000423 | 0.000365 | 0.000471 |
| IRS2 | 1 (fixed) | 0.755684 | 0.28822 | 5.14E-06 | 3.16E-06 | 2.1E-06 |
| Akt | 1.038483 | 0.937216 | 0.897796 | 0.06567 | 0.041611 | 0.036028 |
| GSK3β | 1.248498 | 1.233192 | 1.348061 | 0.15389 | 0.10087 | 0.096531 |
| FoxO1 | 1.476479 | 1.324451 | 1.104076 | 0.166243 | 0.098556 | 0.071852 |
| ACKY | 1.346885 | 1.232721 | 0.615567 | 0.200196 | 0.122785 | 0.053806 |
| mTor | 1 (fixed) | 1 (fixed) | 1 (fixed) | 0.000195 | 0.000124 | 0.000107 |
| X | 1 (fixed) | 1 (fixed) | 1 (fixed) | 0.104084 | 0.06857 | 0.059923 |
| S6K | 58.61192 | 58.36148 | 54.26853 | 0.170797 | 0.163601 | 0.150732 |
| Y | 1 (fixed) | 1 (fixed) | 1 (fixed) | 8.46E-06 | 5.36E-06 | 4.64E-06 |
| 4E-BP1 | 1 (fixed) | 0.962231 | 0.967315 | 0.488727 | 0.449567 | 0.448022 |

